# Effect of reduced graphene oxide (rGO) compaction degree and concentration on rGO-polymer composites printability and cell interactions

**DOI:** 10.1101/2021.05.18.444680

**Authors:** María Cámara-Torres, Ravi Sinha, Siamak Eqtesadi, Rune Wendelbo, Marco Scatto, Paolo Scopece, Alberto Sanchez, Sara Villanueva, Ainhoa Egizabal, Noelia Álvarez, Alessandro Patelli, Carlos Mota, Lorenzo Moroni

## Abstract

Graphene derivatives combined with polymers have attracted enormous attention for bone tissue engineering applications. Among others, reduced graphene oxide (rGO) is one of the preferred graphene-based fillers for the preparation of composites via melt compounding, and their further processing into 3D scaffolds, due to its established large-scale production method, thermal stability, and electrical conductivity. In this study, rGO (low bulk density 10g/L) was compacted by densification using a solvent (either acetone or water) prior to melt compounding, to simplify its handling and dosing into a twin-screw extrusion system. The effects of rGO bulk density (medium and high), densification solvent, and rGO concentration (3, 10 and 15% in weight) on rGO dispersion within the composite, electrical conductivity, printability and cell-material interactions were studied. High bulk density rGO (90 g/L) occupied a low volume fraction within polymer composites, offering poor electrical properties but a reproducible printability up to 15 wt% rGO. On the other hand, the volume fraction within the composites of medium bulk density rGO (50 g/L) was higher for a given concentration, enhancing rGO particle interactions and leading to enhanced electrical conductivity, but compromising the printability window. For a given bulk density (50 g/L), rGO densified in water was more compacted and offered poorer dispersability within the polymer than rGO densified in acetone, and resulted in scaffolds with poor layer bonding or even lack of printability at high rGO percentages. A balance in printability and electrical properties was obtained for composites with medium bulk density rGO densified in acetone. Here, increasing rGO concentration led to more hydrophilic composites with a noticeable increase in protein adsorption. Moreover, scaffolds prepared with such composites presented antimicrobial properties even at low rGO contents (3 wt%). In addition, the viability and proliferation of human mesenchymal stromal cells (hMSCs) was maintained on scaffolds with up to 15% rGO and with enhanced osteogenic differentiation on 3% rGO scaffolds.

## 1. Introduction

Since the isolation of graphene, graphene based materials have been thoroughly exploited for various biomedical applications, and in particular for bone tissue engineering, due to their excellent mechanical, chemical and electrical properties, as well as to their unique ability of directing stem cell differentiation. [1–3] Pristine graphene is a one atom thick, hexagonal lattice structure of sp2 hybridized carbon atoms, which confer all the aforementioned properties. [4–6] Due to high synthesis costs and small scale production of graphene through bottom-up processes, such as chemical vapor deposition, [7] or by direct exfoliation of graphite, such as micromechanical cleavage or direct sonication, [8] currently the most promising method for large scale production of graphene-like materials relies on graphene oxide (GO) as starting material. [9] GO can be easily mass-produced by oxidation of graphite through well-established methods, and used as-prepared or after exfoliation into graphene-like sheets. [9] In addition, GO possesses oxygen containing functional groups disrupting each carbon plane, including epoxy, carbonyl, ketone and hydroxyl groups, which allows for good water and polymers dispersability and offers the possibility of functionalization with biomolecules to tune its bioactivity, unlike graphene sheets directly obtained from graphite. [10] However, GO is electrically insulating and thermally unstable, requiring at least partial reduction to restore these properties when required for the final application. GO can be reduced by different chemical and thermal procedures, leading to exfoliated wrinkled reduced graphene oxide (rGO) sheets, similar to pristine graphene, but with some oxygen defects and holes on the carbon skeleton. [11]

In recent years, graphene, GO and rGO have been used as fillers in both natural and synthetic polymer nanocomposites to improve their physicochemical properties for bone tissue engineering applications. Among different types of such composites, including electrospun fibers, [12, 13] hydrogels [14, 15], or additive manufactured (AM) scaffolds, [16–19] the latter are superior when aiming towards load bearing applications, as they provide higher mechanical properties. Compared to other AM methods, melt extrusion AM (ME-AM) is considered a cost effective and established technique within the tissue engineering field, which enables the processing of a wide range of biocompatible and biodegradable thermoplastic materials, and provides full control on the internal pore architecture of the scaffolds. Accordingly, three main routes have been explored for the production of graphene derivatives-polymer composites to be used by ME-AM: solution blending, melt blending, and *in situ* polymerization. [20] While solvent blending is a simple path to obtain a good filler dispersion, it generally requires the use of expensive and non-environmentally friendly organic solvents, which can potentially stay as residues within the composite matrix. [21] *In situ* polymerization has shown to offer even a higher level of dispersion due to monomer intercalation between filler layers, followed by polymerization. However, this process can also require the use of solvents and is limited to specific polymer types. [22] In spite of not providing the same level of filler dispersion as the aforementioned techniques, melt blending by twin screw compounding holds the most promise for large-scale composite fabrication, due to its lower cost, green production and industrial applicability. [23, 24] However, some reports have described the formation of air pockets within extruded polymer composite filaments containing GO at high concentrations, due to its thermal instability and *in situ* thermal reduction at the extruding temperature. [25] rGO has also offered some challenges when used as a filler, despite its thermal stability. This is due to its very low bulk density and volatility after volume expansion upon thermal reduction, which impedes its free-flowability into melt compounders and can lead to nanoparticle intake by inhalation during handling. [26] To overcome this, rGO compaction or densification by dispersion in a solvent and drying, [24] pre-coating with polymer particles in solution, [27] or the preparation of a highly concentrated masterbatch by solvent blending, prior to melt blending, have been considered. [28] Yet, the attention from rGO as filler for ME-AM scaffolds for bone tissue engineering applications has been deviated and, to the best of our knowledge, the vast majority of previous studies have focused only on graphene and GO fillers for this application, despite pristine graphene’s low yield production and GO’s poor thermal stability and lack of electrical conductivity. [17, 29–31]

Here, we prepared densified rGO and studied, for the first time, the effect of rGO densification on its dispersion within melt blended rGO- poly(ethylene oxide terephthalate)/poly(butylene terephthalate) (PEOT/PBT) composites, at various rGO concentrations. While most of previously reported graphene based ME-AM scaffolds contain only up to 3 wt% filler content, and very rarely up to 10 wt%, in this study we investigated the feasibility of preparing composites with up to 15 wt% rGO, and evaluated their conductivity and printability as a function of rGO compaction and concentration. Moreover, the effect of rGO concentration on the material physicochemical properties, in terms of hydrophilicity, protein adsorption, and antimicrobial properties was assessed. To evaluate its application for bone tissue engineering, we further assessed human mesenchymal stromal cells (hMSCs) adhesion, proliferation and osteogenic differentiation on the prepared 3D ME-AM scaffolds.

## 2. MATERIALS AND METHODS

### 2.1. rGO synthesis and characterization

GO was synthesized by Abalonix AS according to a proprietary in-house GO manufacturing method, and thermally reduced to obtain partially reduced rGO. During the reduction process, GO was introduced for few seconds into a tubular oven at 600 °C. However, due to cold air flowing through the oven chamber, the real temperature that the material experiences was estimated to be ~ 300 °C. Three different rGO batches were produced following the same protocol. Densified / compacted (the two terms will be used interchangeably) forms of these rGOs (hereafter referred to as d-rGO) were produced by dispersion into a solvent and subsequent drying. rGO was dispersed in acetone at 50 mg/ml and dried at room temperature to obtain d-rGO-B, with a bulk density of ~ 50 g/L. To obtain d-rGO-A, rGO was dispersed in water at 50 mg/ml and dried at 100 °C. The water dispersion and drying process was carried out two times to obtain d-rGO-C with a bulk density of ~ 90g/L. Overall, three different d-rGO materials were obtained: d-rGO-A (densified in water, bulk density 50 g/L), d-rGO-B (densified in acetone, bulk density 50 g/L) and d-rGO-C (densified in water, bulk density 90 g/L)

In order to evaluate the atomic composition of each d-rGO material, X-ray photoelectron spectroscopy (XPS, K-Alpha – Thermo Scientific, US) was used. Moreover, X-ray diffraction (XRD, D8 Advanced, Brukers, Germany) was employed to investigate the different rGO batches crystallinity and their layered structure. In addition, SEM was carried out to characterize the microstructure of the different d-rGO materials.

### 2.2. Composite production and characterization

d-rGO-polymer composites were obtained by compounding individually the various d-rGO materials (d-rGO-A, d-rGO-B and d-rGO-C) with PEOT/PBT (300PEOT55PBT45, pellets form, PEO Mw 300 kDa, PEOT:PBT weight ratio= 55:45, intrinsic viscosity 0.51 dl/g, Polyvation, The Netherlands). Production was carried out in a lab scale co-rotating twin-screw extruder installed in Nadir S.r.l., consisting of a screw profile with 8 zones, 3 interposed kneading sections, and a screw dimensions of 11 mm diameter and 40 length-to-diameter ratio. PEOT/PBT pellets and d-rGO powder were fed at different concentrations (3, 10, 15 wt%) into the main hopper with a volumetric feeder. The screw rotation speed was fixed at 80 rpm, while the barrel temperature was set at 140°C for the first zone and 145-150°C for the following zones. Resulting composite wires were taken at the die exit, solidified in air and pelletized in a pelletizer machine. These will be referred to as “X% d-rGO-Y” pellets or composites, where “X” refers to the rGO concentration in wt% (3, 10, 15) and “Y” refers to the d-rGO batch (A, B, or C).

TGA measurements of the d-rGO batches and composites were carried out in a vertical thermo-balance, TGA Discovery 55 analyser (Water-TA Instruments). About 11 mg samples were conducted under nitrogen atmosphere (99,999% N_2_) with a flow rate of 90 ml/min and using a temperature heating rate of 10 °C/min up to 900 °C. Residual weight percentages at 550 °C extracted from the TGA curves were used to calculate the experimental rGO loading of each composite. The 550 °C temperature was selected from the separate rGO and polymer only TGA’s, as the temperature where most of the polymer was burnt off and most of the rGO was still left.

To analyze potential aggregation of d-rGO after the compounding process, pellets of each material were dissolved in chloroform. After mechanical stirring, a sample of each solution was dispensed on a scanning electron microscopy (SEM) sample holder, dried at room temperature, and imaged using SEM (XL-30) operating at a beam voltage of 15 kV and a spot size of 3. Particle size distribution was assessed using the image analysis software ImageJ.

### 2.3. Scaffolds fabrication and characterization

Scaffolds were fabricated via ME-AM. The platform consisted of a custom-made printhead, with separate heating sources for the cartridge (polymer reservoir) and extrusion screw, mounted on a three-axis stage (Bioscaffolder, Gesim). [32] Briefly, the cartridge was filled with the pellets of the composite material to be printed, and scaffolds with a 0-90 architecture, 250 μm fiber diameter, 200 μm layer thickness and 750 μm strand distance (center to center) were fabricated according to parameters in Table 1. Cylindrical scaffolds of 4 mm diameter and 4 mm height were punched out from 15×15×4 mm^3^ manufactured blocks using a biopsy punch and used for further experiments.

**Table 1.** Fabrication parameters of d-rGO-polymer composite scaffolds

| Scaffold | T reservoir | T screw | Pressure | Screw rotation | Translation |
| --- | --- | --- | --- | --- | --- |

|  | (° C) | (° C) | cartridge<br>(bar) | speed (rpm) | speed (mm/s) |
| --- | --- | --- | --- | --- | --- |
| **3% d-rGO-A** | 195 | 200 | 8 | 60 | 15 |
| **10% d-rGO-A** | 220 | 220 | 8 | 60 | 15-20 |
| **15% d-rGO-A** |  |  | not extrudable |  |  |
| **3% d-rGO-B** | 195 | 200 | 8 | 60 | 15 |
| **10% d-rGO-B** | 200 | 210 | 8 | 40 | 15-20 |
| **15% d-rGO-B** | 200 | 210 | 8 | 60 | 15-20 |
| **3% d-rGO-C** | 200 | 205 | 8 | 60 | 15 |
| **10% d-rGO-C** | 200 | 205 | 8 | 60 | 15 |
| **15% d-rGO-C** | 200 | 210 | 8 | 60 | 15-20 |

Scaffolds morphology and porosity were assessed using a stereomicroscope (Nikon SMZ25). Scaffolds surface roughness was examined using SEM, operating at 10 kV, and a spot size of 3. The electrical conductivity of extruded filaments (length 6 cm, diameter 340 μm and 800 μm) of each material was measured using a digital multimeter (maximum measurable resistance: 2000 MΩ). To do this, filaments with a known diameter and length were connected to the multimeter electrodes with conductive silver paste (Chemtronics Silver Conductive Adhesive Epoxy) to ensure good contact. The volume electrical conductivity was calculated by Equation 1.

*Equation 1. Volume electrical conductivity (*σ*) formula, where l is the filament length (6 cm), R is the electrical resistance and A is the filament cross sectional area (diameter 340 μm and 850 μm).*

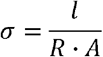

Cell studies and antimicrobial analysis was performed only on the d-rGO-B scaffolds due to their superior combined properties for the final application, in terms of processability and electrical conductivity, compared to the d-rGO-A and d-rGO-C composites.

### 2.4. Films preparation and characterization

2D films were prepared from around 60 milligrams of (3, 10, 15)% rGO-B pellets and PEOT/PBT pellets, which were molten at 190-210 °C and pressed with a coverslip against a Teflon sheet to obtain 14 mm diameter, ~300 μm thickness films. Static contact angle was measured on these films using the sessile drop method. To do this, a 4 μl water droplet was placed on top of the films by an automatic syringe dispenser (Krüss DSA25S). 20 seconds after droplet formation, the contact angle was calculated automatically by the device’s software using the Laplace–Young curve fitting.

Protein adsorption to the films was analyzed by incubation in a bovine serum albumin-FITC protein solution (1mg/ml, Sigma Aldrich) overnight at 37 °C. After washing in PBS, films were blotted in an adsorbent paper and incubated for 2h at RT in a 1% SDS solution. Supernatants fluorescence was measured at excitation/emission = 495 nm/519 nm. Cell studies and antimicrobial analysis was performed only on the d-rGO-B films, due to their superior combined properties for the final application, in terms of processability and electrical conductivity, compared to the d-rGO-A and d-rGO-C composites.

### 2.5. Antibacterial activity

The antibacterial activity of (3, 10, 15)% d-rGO-B scaffolds and films against *P. aeruginosa* (CECT 116) and *S. epidermidis* (CECT 231) was evaluated by means of the flask shake test method. Briefly, scaffolds were immersed in concentrated bacterial suspensions (10^6^ CFU·L^−1^) in nutrient broth (1:500) for 24 h at 37 °C. After incubation, the number of viable bacteria present in the suspensions was measured by placing aliquots of the suspensions in sterile petri dishes with molten nutrient agar per triplicate and swirled gently. The petri dishes were incubated at 37 °C for 24 h, after which the colonies present on the plates were counted. Values are reported as the log base 10 reduction (R), calculated as the difference in the log base 10 of the viable cell counts found on a suspension that has not been in contact with the scaffolds and the counts on a suspension that has been in contact with the scaffolds.

In addition, the antibacterial activity of (3, 10, 15)% d-rGO-B scaffolds was evaluated by means of the Agar disk-diffusion method. For that, scaffolds were incubated for 3 days in 1 ml PBS. Mueller-Hinton agar plates were spread with a standardized inoculum of *P. aeruginosa* and *S. epidermidis*. At each timepoint (24h, 48h, 72h), 20 μl of each of the scaffolds’ supernatant was collected and used to impregnate commercial paper discs, which were subsequently placed on the agar surface. The agar plates were incubated under 37 °C during 18-24 hours. After incubation, zones of growth inhibition (ZOI) around each of the discs (including disc diameter) were measured to the nearest millimeter. For reporting, the disc diameter was subtracted.

### 2.6. Cell seeding and culture

HMSCs isolated from bone marrow were purchased from Texas A&M Health Science Center, College of Medicine, Institute for Regenerative Medicine (Donor d8011L, female, age 22). Cryopreserved vials at passage 3 or 4 were plated at a density of 1000 cells×cm^−2^ in tissue culture flasks and expanded until approximately 80 % confluency in complete media (CM). CM was composed of αMEM with Glutamax and no nucleosides (Gibco) supplemented with 10% fetal bovine serum (FBS), without penicillin-streptomycin (PenStrep) at 37 °C and 5% CO_2_.

#### 2.6.1. Cell seeding on 2D films

PEOT/PBT and (3, 10, 15)% d-rGO-B films were disinfected in 70% ethanol for 20 min, washed 3 times with PBS, and fixed on the bottom of well-plates with the help of biocompatible O-rings (Eriks, 10023241). Prior to cell seeding, films were incubated in CM for 2h at 37 °C and 5% CO_2_ for protein adhesion. Trypsinized hMSCs were re-suspended in fresh media and seeded at 2500 cells per film. HMSCs were cultured on films for 7 days, and media was replaced every 2 or 3 days.

#### 2.6.2. Cell seeding on scaffolds

For cell attachment experiments, PEOT/PBT and (3, 10, 15)% d-rGO-B scaffolds were disinfected in 70% ethanol and incubated in CM for 2h at 37 °C and 5% CO_2_ to allow protein attachment. Subsequently, scaffolds were blotted on top of a sterile filter paper and placed in the wells of a non-treated wellplate. Trypsinized hMSCs were resuspended in a dextran solution (500 kDa, Farmacosmos) (10 wt% dextran in CM), to achieve uniform cell distribution, [33] and were seeded at a density of 2×10^5^ cells with 37 μl of CM per scaffold. After 4h incubation for cell attachment, scaffolds were transferred to new wells containing 1.5 ml of basic media (BM) (CM supplemented with 200 μм L-Ascorbic acid 2-phosphate). BM was replaced after 24h and every 2 or 3 days from then on. Scaffolds were cultured for 7 days. For osteogenic differentiation experiments, PEOT/PBT and 3%d-rGO-B scaffolds were seeded as mentioned before, and cultured for 7 days in BM, after which scaffolds were further cultured either in BM or in mineralization media (MM), consisting of BM supplemented with dexamethasone (10 nм) (Sigma-Aldrich) and β-glycerophosphate (10 mм) (Sigma-Aldrich) for another 28 days.

#### 2.6.3. Cells imaging on films and scaffolds

Films and scaffolds seeded and cultured with cells were fixed in 4% paraformaldehyde and incubated for 30 min in Triton-X 100 (0.1 v%). Then, cell cytoskeleton and nuclei were stained with 488 Alexa Fluor Phalloidin (Thermo Fisher Scientific, 1:75 dilution in PBS, 1h at RT) and DAPI (0.1 μg/ml in PBS, 15 min), respectively.

#### 2.6.4. Biochemical assays

##### 2.6.4.1. Alkaline Phosphatase (ALP) activity

To measure ALP activity, scaffolds were collected after 14 and 35 days of culture, freeze-thawed 3 times and incubated for 1h at RT in a cell lysis buffer composed of KH_2_PO (0.1 м), K_2_HPO_4_ (0.1 м), and Triton X-100 (0.1 v%), at pH 7.8. The chemiluminescent substrate for alkaline phosphatase CDP star® ready to use reagent (Roche) was added to the cell lysate at a 1:4 ratio, and luminescence was measured using a spectrophotometer. Remaining cell lysates were kept for DNA quantification. ALP values were reported normalized to DNA content.

##### 2.6.4.2. DNA assay

DNA assay was performed on cells cultured on films and scaffolds using the CyQUANT cell proliferation assay kit (Thermo Fisher Scientific). Lysed scaffolds after ALP assay and frozen films after collection were incubated overnight at 56 °C in Proteinase K solution (1 mg×ml^−1^ Proteinase K (Sigma-Aldrich) in Tris/EDTA buffer) for matrix degradation and cell lysis. After three freeze-thawing cycles, samples were incubated for 1h at RT with a 20X diluted lysis buffer from the kit containing RNase A (1:500) to degrade cellular RNA. Subsequently, the fluorescent dye provided by the kit (1:1) was added and fluorescence was measured after 15 min incubation (emission/excitation = 520/480 nm). DNA concentrations were calculated from a DNA standard curve.

#### 2.6.5. Alizarin red staining and quantification

Calcium deposits on scaffolds cultured for 35 days in BM and MM were stained by alizarin red S (ARS) (60 mм, pH 4.1 - 4.3, 20 min incubation at RT) and visualized using a stereomicroscope (Nikon SMZ25). After imaging, ARS was extracted and quantified. To do this, scaffolds were incubated for 1h at RT with 30 v% acetic acid, followed by 10 min incubation at 85 °C. After a centrifugation step at 20,000 rcf for 10 min, ammonium hydroxide (5 м) was added to the supernatants to bring the pH to 4.2. The absorbance was measured at 405 nm using a spectrophotometer. Concentration of ARS was calculated from an alizarin red standard curve and the values were normalized to DNA content.

## 3. RESULTS AND DISCUSSION

### 3.1. Effect of densification on resulting d-rGO properties

One of the most attractive methods for dispersing fillers into polymers, in terms of costs, scalability and green production, is melt compounding. [34] Due to the low scale separation of exfoliated graphene sheets from bulk graphite, and the low thermal stability of GO, the use of rGO within melt blended polymer-graphene derivatives composites is preferred. [35] However, the low bulk density of rGO makes its feeding into compounders very difficult. This low bulk density originates from the exfoliation and volume expansion of GO caused by the CO_2_ formed upon the decomposition of oxygen-containing functional groups during the high temperature thermal reduction process. [24, 26, 28] For this reason, prior to compounding with PEOT/PBT, the as-produced very low bulk density rGO (10 g/L) was densified (d-rGO) to ease its handling and dosing into the twin-screw compounding system, as well as to reduce the nanoparticles’ inhalation hazard. In order to investigate the effect of different densification parameters (i.e. densification solvent and final rGO bulk density) on the resulting d-rGO properties, as well as on the compounding and printing processes, two different densification solvents, acetone and water, were used. Previous literature suggests the applicability of these solvents as densification medium. [24, 36][37] Here, three d-rGO materials were obtained after dispersion in either of these two solvents (Figure 1): i) d-rGO-A: densified in water, bulk density 50 g/L (ρ _medium_), ii) d-rGO-B: densified in acetone, bulk density 50 g/L (ρ _medium_), and iii) d-rGO-C: densified in water, bulk density 90 g/L (ρ _high_).

**Figure 1.**
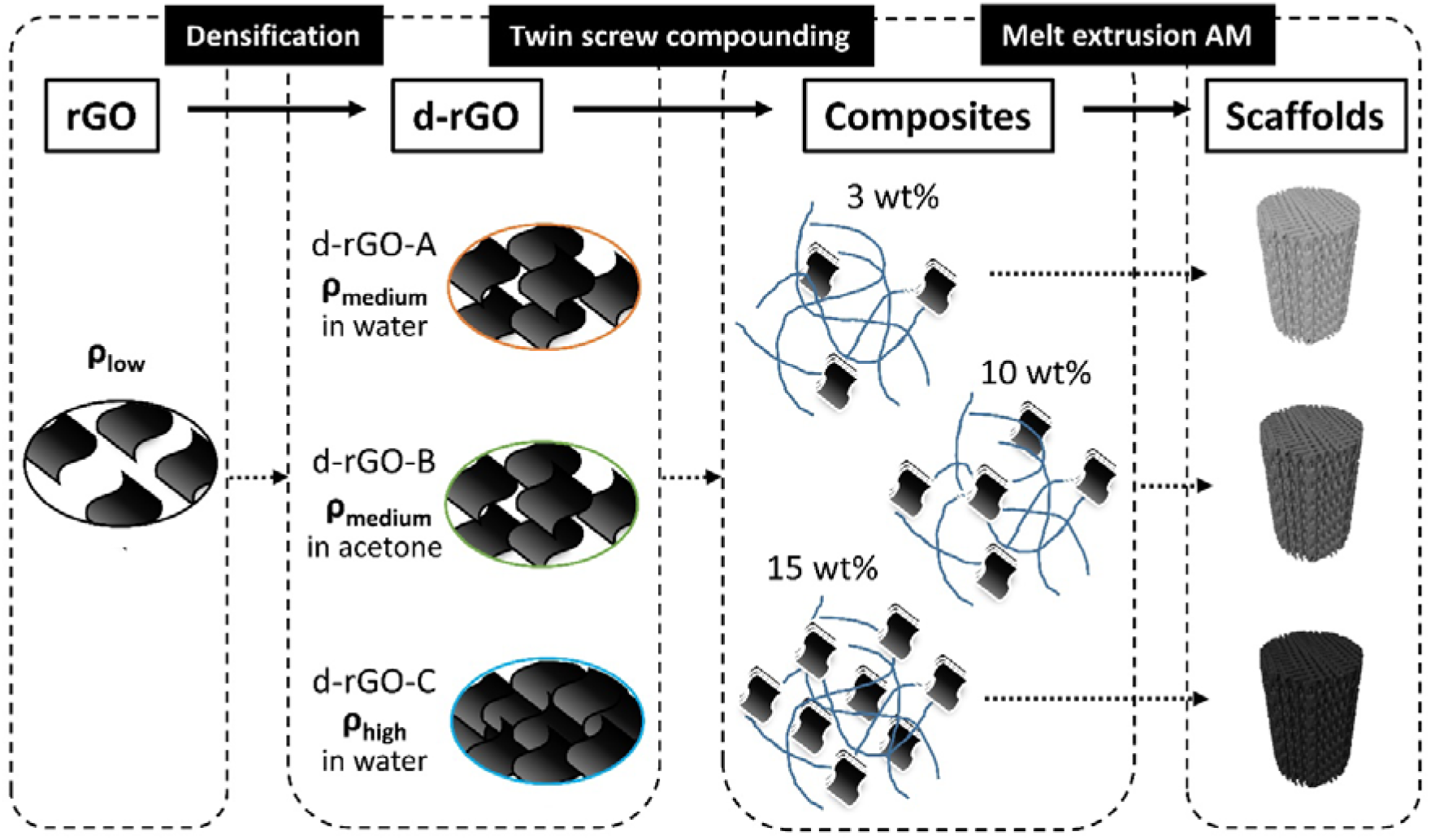
Schematic of the polymer-rGO composite scaffold fabrication route. The low bulk density as-prepared rGO was densified (d-rGO), and three different d-rGO types were obtained: d-rGO-A (densified in acetone, medium bulk density), d-rGO-B (densified in water, medium bulk density) and d-rGO-C (densified in water, high bulk density). Each of the d-rGO was melt blended via twin screw extrusion with PEOT/PBT to obtain composites at three different d-rGO concentrations (3, 10 and 15 wt%). Each of the composites was used to fabricate scaffolds via melt extrusion AM.

Since each of the d-rGO came from a different rGO source, their elemental composition was analysed to confirm equal C/O ratios and to allow for subsequent comparisons. According to the XPS analysis in Table S1, all d-rGO materials were partially reduced, due to the presence of oxygen in their composition. The C/O ratios were maintained between 5 and 6 in all rGO batches, verifying a consistent reduction degree among batches. Moreover, all batches presented traces of N, Al, Si, S, Cl or Fe lower than 1 atomic%. Impurities of sulfur and nitrogen were likely to be due to the covalently bonded sulfates and nitrates during the Hummers method to produce GO. [38, 39] Metallic contaminations are thought to derive from the starting graphitic material itself or the synthesis process. [40, 41] The XRD patterns also showed no differences in the crystalline structure among the different d-rGO’s (Figure S1). All diffractograms showed a (002) diffraction peak at 24 degrees, corresponding to rGO materials with an interlayer distance of 0.37 nm, and a (100) peak at ~ 42 degrees. [38, 42] Upon calculation of the full width at half maximum (FWHM) of the (002) peaks, the same crystallite size among the different d-rGO’s was confirmed (Figure S1B). The low intensity and broadness this peak revealed the amorphous structure and the short-range order of the d-rGO’s. [43, 44] Other peaks at ~28-33 degrees, which correspond to crystalline mineral impurities, were also appreciated in some the diffractograms. [45] In addition to these, some spectra showed low intensity sharp peaks at ~ 26.5 degrees, which correspond to traces of unoxidized graphite. Overall, all d-rGO display similar XRD diffractrograms, suggesting that the densification step did not have an impact on the reduction degree, nanocrystalline structure or crystallite size of the samples. However, it is believed that the high porosity of as-prepared rGO created due to volume expansion and CO_2_ formation during the thermal reduction process, is reduced during densification, leading to a more compacted and aggregated material. rGO aggregation has been previously reported during the complete removal of water after an aqueous GO reduction process, or after drying rGO in an acetone solution. [24, 46] Furthermore, previous studies have revealed that re-dispersion of rGO in water or ethanol followed by a drying step, lead to agglomeration of the powder and a shrinkage of their macroporous and mesoporous structure. [37] For these reasons, we hypothesize that d-rGO-C, with the highest bulk density, was the highest compacted material after solvent evaporation. This was confirmed by SEM, where both d-rGO-B and d-rGO-C displayed particles aggregating together, but d-rGO-C clearly revealed a less porous and more compacted structure at higher magnifications (Figure S2). Although possessing the same bulk density, differences among d-rGO-A and d-rGO-B were also expected, since they were densified in different solvents. Previous literature suggests that the shrinkage of rGO pore size after drying aqueous solutions is more pronounced compared to that of rGO dried in solvents with lower polarity, surface tension, and wettability, such as ethanol or, in our case, acetone. [37] This is due to the better interaction of water with rGO, which exerts a higher capillary force upon evaporation, causing a significant decrease in the macro- and mesopore volume. For this reason, d-rGO-A densified in water is hypothesized to be a more compacted material than d-rGO-B. Therefore, the three different d-rGO could be classified according to their compaction degree, being d-rGO-C the most compacted (densified in water, bulk density 90 g/L), followed by the medium compacted d-rGO-A (densified in water, bulk density 50 g/L), and the low compacted d-rGO-B (densified in acetone, bulk density 50 g/L).

### 3.2. Effect of d-rGO densification on melt compounding

After densification, each of the d-rGO materials was melt compounded with PEOT/PBT to form PEOT/PBT-d-rGO composites (Figure 1). TGA measurements of the experimental d-rGO loading on each of the composites suggested the reproducibility of the melt compounding process regardless the d-rGO used, for all d-rGO concentrations (Table 2, Figure S3). Yet, it is worth noting that when d-rGO-B was blended at high concentrations (15 wt%), the experimental values deviated slightly from the theoretical ones, and the composites contained lower d-rGO amounts than expected. This can be likely due to an inhomogeneous distribution of d-rGO within the pellets and the low amount of sample (a few pellets at most) used for TGA measurements.

**Table 2.** Experimental loadings (wt%) of each of the composites calculated by TGA.

|  | 3 wt% | 10 wt% | 15 wt% |
| --- | --- | --- | --- |
| **d-rGO-A** | $2.91 \pm 0.34$ | $11.22 \pm 0.45$ | $17.73 \pm 0.28$ |
| **d-rGO-B** | $2.18 \pm 0.84$ | $11.68 \pm 1.03$ | $11.73 \pm 1.29$ |
| **d-rGO-C** | $4.64 \pm 0.28$ | $9.47 \pm 6.09$ | $15.01 \pm 1.03$ |

The differences in compaction degree mentioned above correlated well with the filler dispersion within the PEOT/PBT-d-rGO composite after compounding. Filler aggregation as an indirect measure of d-rGO dispersion was evaluated by dissolving the composites in chloroform, and visualizing and quantifying the size of d-rGO particles. Figure 2A displays representative SEM images of d-rGO within each of the composites. Small particles up to 50 μm were abundant for all conditions (relative frequency ~ 80 %) (Figure 2A, B). In addition, big d-rGO aggregates were also observed mainly in d-rGO-A composites (ρ_medium_), at all d-rGO concentrations, and on d-rGO-B composites (ρ_medium_) at high concentrations (10% and 15%). Interestingly, d-rGO-C composites (ρ_high_) displayed much lower d-rGO aggregation, as revealed by the SEM images and the lower number of > 200 μm aggregates compared to d-rGO-A and d-rGO-B composites (Figure 2B). Among all, the d-rGO-A composites displayed the largest number of > 200 μm aggregates, especially at 3% and 15% d-rGO. It is believed that the high bulk density d-rGO-C occupied a smaller volume fraction within the composite at a given wt% compared to d-rGO-A and d-rGO-C, even for high d-rGO concentrations. This avoided particle interaction within the composites and reduced the number of visible aggregates. On the other hand, due to the lower bulk density of d-rGO-A and d-rGO-B, these were able to reach a critical volume occupancy within the composites at much lower concentrations (wt%). This increased the probability of d-rGO particles interaction, and the formation of particles overlaps into bigger agglomerates, by hydrophobic interactions. [10, 47, 48] In addition, due to the lower porosity and higher compaction degree of d-rGO-A compared to d-rGO-B, filler exfoliation and dispersion within the d-rGO-A composites was more difficult, leading to the visualization of a larger number of aggregates. Compared to melt compounding, other composite production methods such as solvent blending or *in situ* polymerization, have shown to ensure more uniform d-rGO dispersions. [24, 49] Moreover, several reports have suggested the preparation by solvent blending of rGO-polymer masterbatches, to be used as starting material to prepare other rGO composites by melt compounding. [28, 50, 51] This approach has shown to lead to better rGO dispersion, as it skips the densification or compaction step. However, it involves the use of organic solvents in the process, which is not desirable for biological applications, as the solvents can remain as residues within the composite matrix.

**Figure 2.**
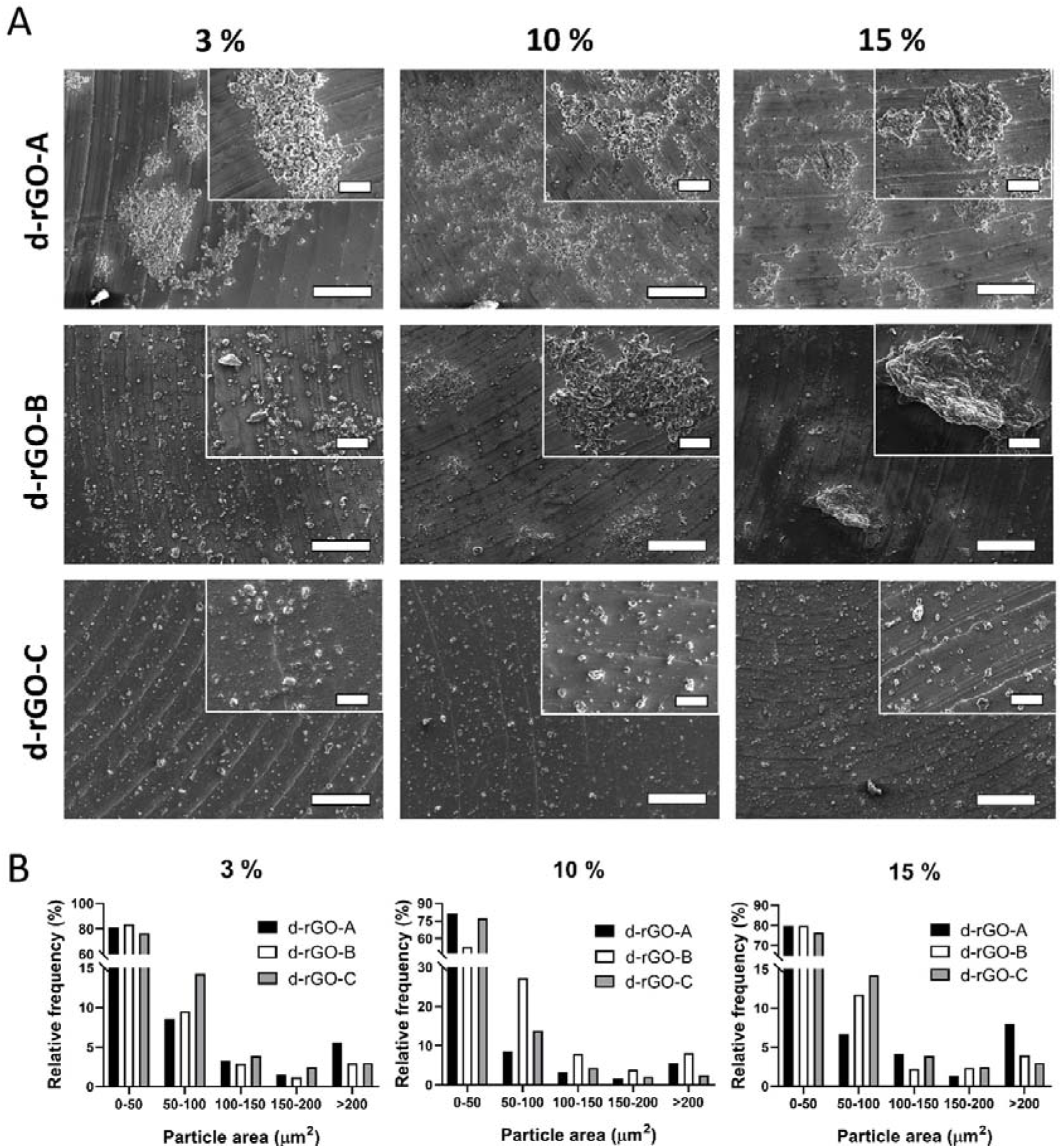
(A) Representative SEM micrographs of d-rGO particles and aggregates after dissolving each composite in chloroform. Scale bars 200 μm. Inserts correspond to high magnification images. Scale bars 50 μm. (B) Size distribution of d-rGO particles and aggregates measured from SEM micrographs in (A).

Notably, d-rGO volume occupancy and dispersion within the composite also correlated well with the electrical properties of extruded composite filaments, as presented in Table 3. d-rGO-C extruded filaments’ high resistance (out of the device range) impeded the report of their conductivity values at any d-rGO concentration. In case of the d-rGO-A and d-rGO-B filaments, their electrical conductivity was measurable from 10% d-rGO, and they were more conductive with increasing d-rGO content, as previously reported. [28] Interestingly, the conductivity of 10% d-rGO-A filaments was 35-fold and 3-fold higher compared to 10% d-rGO-B and 15% d-rGO-B, respectively, suggesting the lower percolation threshold of the d-rGO-A composites. As 15% d-rGO-A was not extrudable, no filament was obtained for this measurement. It is evident that the high volume fraction and poor dispersability of d-rGO-A within the composites helped to form a more connected d-rGO network leading to conductive composites at lower loadings than the less compacted and better dispersed d-rGO-B. On the other hand, the lower volume fraction of d-rGO-C within the composites could not create such conductive pathways and electrical conductivity was not measurable.[24, 46, 52, 53]

**Table 3.** Conductivity values of extruded filaments from each composite.

|  | Conductivity<br>3% (mS/m) | Conductivity<br>10% (mS/m) | Conductivity<br>15% (mS/m) |
| --- | --- | --- | --- |
| **d-rGO-A** | < 0.32 | $13.8 \pm 1.1$ | n.a. |
| **d-rGO-B** | < 0.32 | $0.44 \pm 0.1$ | $5.4 \pm 1.2$ |
| **d-rGO-C** | < 0.32 | < 0.32 | < 0.32 |

### 3.3. Effect of d-rGO compaction on composite printability

The d-rGO volume occupancy, compaction degree and dispersion also influenced composites printability by ME-AM (Figure 3). Interestingly, due to the large volume fraction and big d-rGO aggregates in the 15% d-rGO-A composites, this material was not printable within the operational limits of the AM device (in terms of printing temperature and cartridge pressure), as it remained a paste-like material rather than a melt, even at 220 °C, and a scaffold could not be obtained. Reducing the filler loading down to 3 wt% made it possible to produce scaffolds with a fiber diameter of 250 μm (Figure 3 and Figure S4). The fabrication of 10% d-rGO-A scaffolds was challenging, since lack of bonding between layers prevented their stability and handling, in spite of printing at the highest operational temperature (220 °C) and increasing the fiber diameter (up to 340 μm) for promoting a bigger area of contact in between layers (Figure 3 and Figure S5). On the other hand, scaffolds with a fiber diameter of 250 μm were obtained with all d-rGO-B composites, up to 15% d-rGO-B (Figure 3, Figure S4). Printing temperature was adjusted for each composite, ranging between 195 °C and 210 °C, since viscosity of the melt was found to increase with increasing d-rGO concentration for all composites. [54] Notably, while 3 and 10 % d-rGO-B scaffolds fabrication was reproducible in terms of resulting filament morphology and z-porosity, 15% d-rGO-B scaffolds lateral porosity did not always remain constant across the scaffold height (Figure 3). This was possibly due to sudden viscosity changes during the extrusion process, probably caused by the discontinuous presence of d-rGO-B aggregates changing the flow of the molten composite and, therefore, causing filaments sagging at some points during the printing process. Interestingly, d-rGO-C scaffolds printing was very stable, and scaffolds with optimum lateral porosity were obtained for all d-rGO concentrations (Figure 3). Looking at d-rGO-C scaffolds filament cross section by SEM (Figure S6), it can be observed that PEOT/PBT (showing a flowy appearance) occupies much larger volume than d-rGO particles (with a grainy and spiky appearance), which are present as dispersed islands within a polymer matrix for all d-rGO concentrations, making these materials relatively easy to print. On the other hand, d-rGO dominantly occupied the d-rGO-A and d-rGO-B scaffolds filaments volume, especially on 10% and 15% d-rGO scaffolds, leading to difficulties when printing. Overall, these results suggest that d-rGO bulk density, and therefore volume fraction, as well as d-rGO densification solvent, which dictates compaction and dispersability degree, plays key roles in the maximum filler loading to process composites into ME-AM scaffolds. While previous studies have reported ME-AM scaffolds production only up to 10 wt% pristine graphene or rGO, [19, 31] we were able to produce scaffolds with up to 15 wt% d-rGO. Despite the apparent lower rGO content in these studies, equivalent volume percentages or graphene densities would be required to make fair comparisons with the composites studied herein. Nevertheless, to the best of our knowledge, solvent blending and direct ink writing (solvent-based printing) are the only described manufacturing methods for scaffolds with higher graphene derivatives concentrations (50-75 wt% graphene), which both suffer from requiring the application of organic solvents. [18, 53]

**Figure 3.**
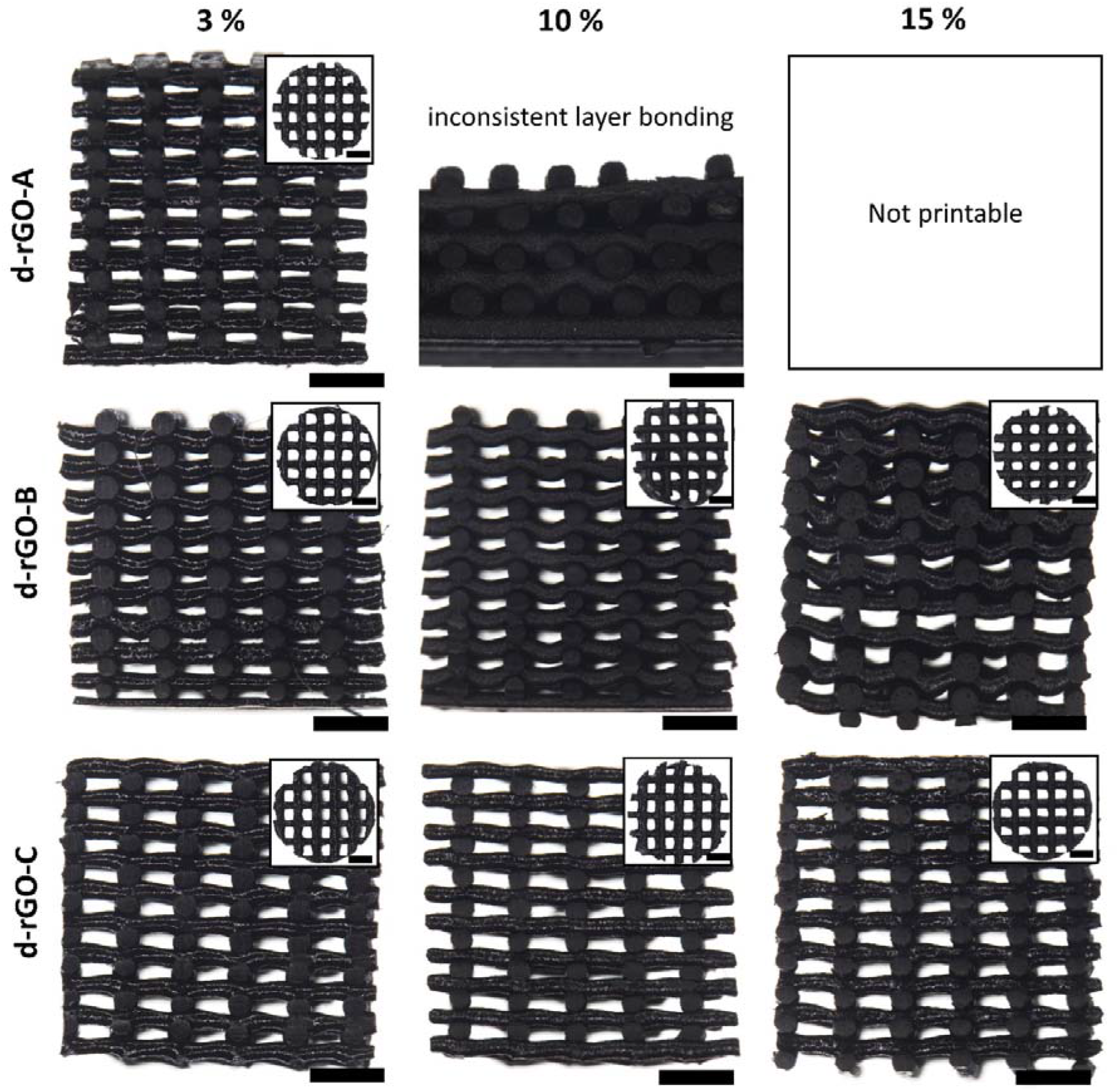
Stereomicroscopy images of 3D ME-AM scaffolds cross sections and top view (inserts) obtained using each of the d-rGO composites, depicting scaffolds morphology and interconnected porosity. Scale bars 1mm.

The ME-AM scaffolds microscale surface roughness was analysed by SEM (Figure 4). As a general observation, the qualitative scaffold surface roughness increased with increasing d-rGO content for all scaffolds types, given by the irregular protuberances formed by the underlying rGO particles. Interestingly, the surface of 3% d-rGO scaffolds remained rather smooth, regardless of the d-rGO type (Figure 4). This is in agreement with previous reports, in which the surface of ME-AM scaffolds containing less than 5 wt% graphene (or derivatives) did not present changes in their surface microroughness compared with bare polymeric scaffolds. [55] In the case of 10% and 15% d-rGO scaffolds, the degree of roughness was influenced by the type of d-rGO, i.e. its bulk density, compaction degree and volume fraction within the filament, for a given d-rGO concentration. In this regard, 10% d-rGO-A presented higher roughness than 10% d-rGO-B, and both higher than 10% d-rGO-C. Similarly, 15% d-rGO-B scaffold surface was much rougher than that of 15% d-rGO-C scaffolds, where only few wrinkles raised on the surface. This is likely due to the much lower volume fraction occupied by the highly compacted d-rGO-C particles decreasing their probability of populating the filaments surface.

**Figure 4.**
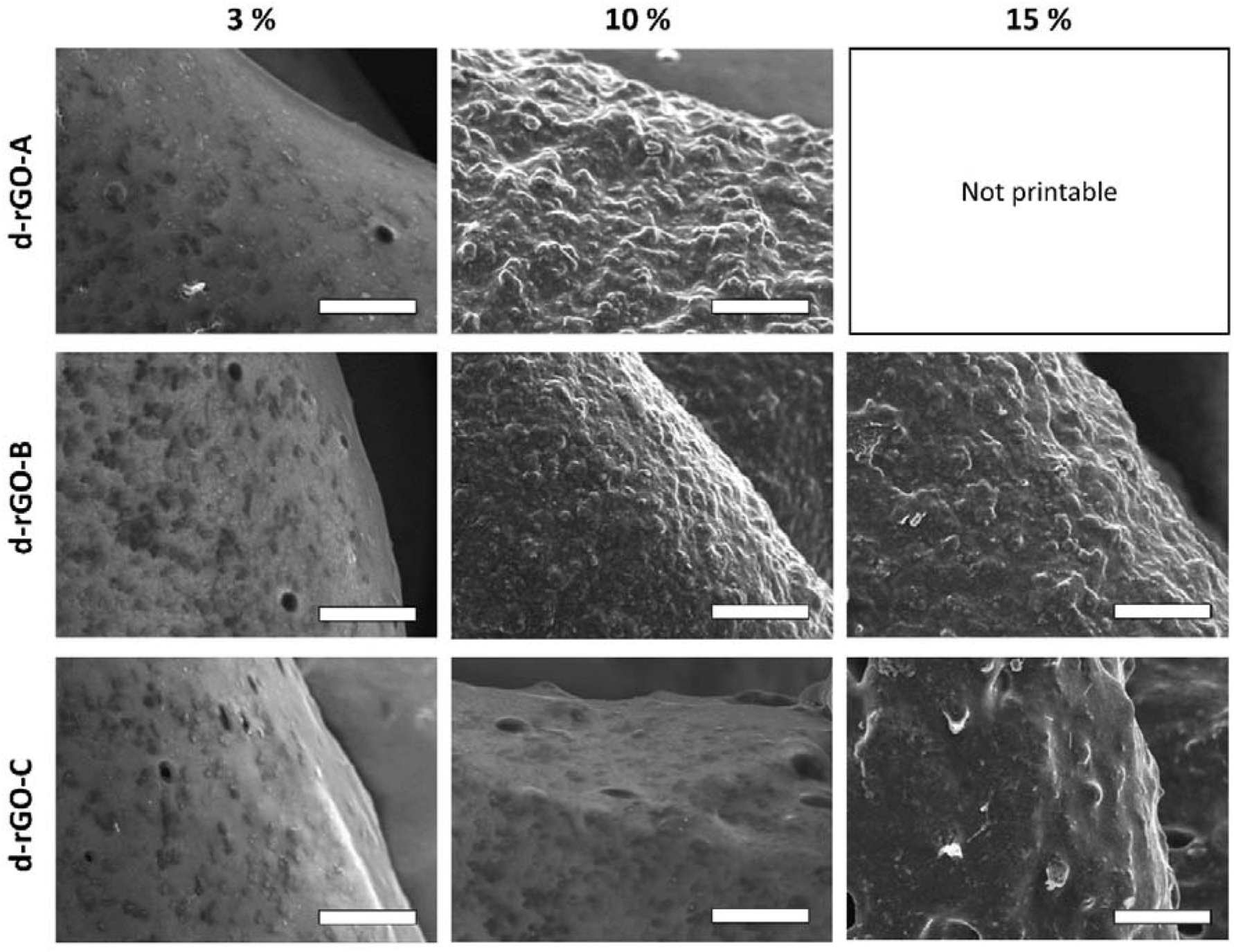
Representative SEM micrographs of composite scaffolds depicting the scaffold’s surface roughness. Scale bars 50 μm.

### 3.4. Effect of d-rGO concentration on wettability and protein adsorption

Prior to assessing the interaction of cells with the ME-AM scaffolds, different material properties, such as wettability and protein adsorption, as well as cell-material interactions in 2D were evaluated. For this and further studies, only the d-rGO-B composites were taken into consideration as they demonstrated a wider printability window (up to 15 wt%) compared to d-rGO-A composites, and superior electrical conductivity compared to the d-rGO-C composites, properties that are considered relevant for their application in bone tissue regeneration. The static contact angle measurements on composite films (Figure 5A), suggested that the material remained rather hydrophobic (contact angle ~80 °) up to 10% d-rGO-B. 15% d-rGO films contact angle was reduced down to ~50 °, denoting the composite hydrophilicity. While the increased hydrophilicity of polymers upon d-rGO-B addition is in agreement with some previous reports, [19, 56] others have suggested the opposite trend. [57, 58] The increase in hydophilicity of 15% d-rGO-B observed within our study might be explained by the d-rGO-B being partially reduced and containing remnant hydrophilic oxygen functional groups, as well as by the presence of larger amounts of d-rGO on the surface of the films with increasing d-rGO-B concentrations. Protein adsorption was assessed by incubating the films in a BSA solution (Figure 5B). While 3% d-rGO-B films adsorbed as much protein as PEOT/PBT films, d-rGO-B content over 3% led to a significant increase in protein adsorption to the films. It is believed that rGO interacts with proteins mainly through hydrophobic van der Waals and π−π stacking interactions, due to the carbon structure. [59, 60] Yet, electrostatic interactions and hydrogen bonding are also possible due to the partial presence of oxygen groups on rGO. [59, 60] As Figure 5B demonstrates, increasing the d-rGO-B concentration up to 15% led to a 4-fold increase in protein adsorption in 15% d-rGO-B films, likely due to a larger amount of interaction points with proteins, compared to lower d-rGO-B content films.

**Figure 5.**
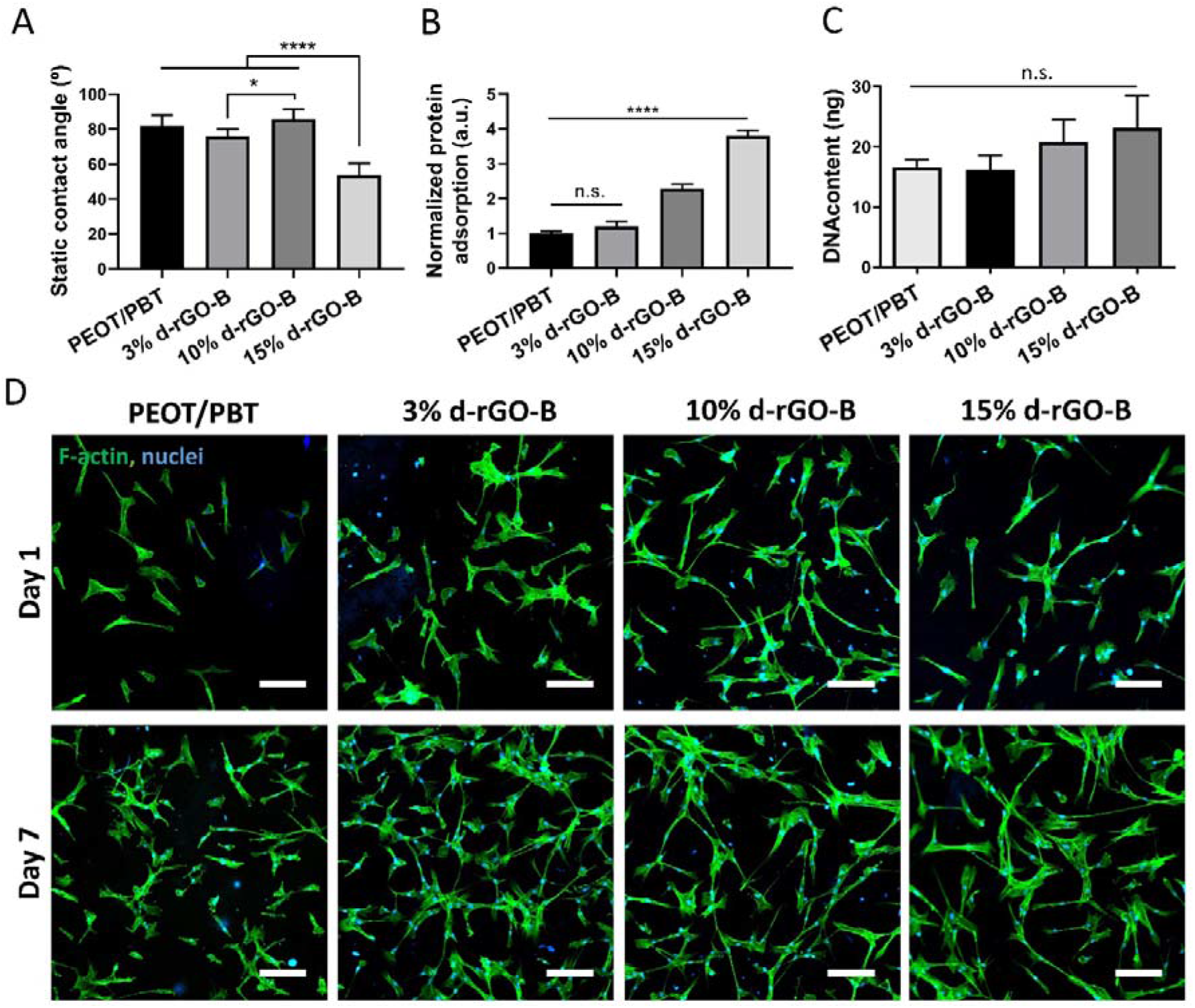
Effect of rGO concentration on physicochemical properties and cell-material interactions in 2D. (A) Static contact angle measured on PEOT/PBT and d-rGO-B composite films. (B) Protein adsorption to d-rGO-B films upon incubation in a BSA solution, normalized to protein adsorption to PEOT/PBT films. (C) Quantification of hMSCs attachment to d-rGO-B films after 1 day of culture. (D) Fluorescent microscopy images of hMSCs (F-actin in green, nuclei in blue) cultured on films for 1 and 7 days. Scale bars 100 μm. Data presented as average ± s.d. and statistical significance performed using one-way ANOVA with Tukey’s multiple comparison test (n.s. p > 0.05, * p < 0.05, **** p < 0.0001).

Surface physicochemical properties, i.e. roughness, hydrophilicity and protein adsorption ability, can regulate cellular behaviour. To evaluate this effect, hMSCs were cultured on films pre-incubated in cell culture medium. A slight increase, yet not significant, in cell attachment with increasing d-rGO-B content was observed upon DNA quantification at day 1 (Figure 5C). Compared to the spread morphology of cells on PEOT/PBT films, hMSCs cultured on d-rGO-B based films, especially at high d-rGO-B content (10 and 15 wt%), showed a spindle-shape, elongated and branched morphology at both day 1 and 7, suggesting higher affinity of hMSCs towards d-rGO-B containing materials, probably due to the aforementioned enhanced protein adsorption (Figure 5D). It is also plausible that the ridges and groves on the films rough surface, given by the underlying d-rGO-B, provided contact guidance cues for cells to acquire an elongated morphology. [61, 62]

### 3.5. Effect of d-rGO concentration on scaffolds antimicrobial activity

The antimicrobial activity of scaffolds was evaluated against relevant Gram - and Gram + bacterial strains in the orthopedic field: *P. aeruginosa* and *S.epidermidis* respectively. Preliminary experiments demonstrated the biocidal activity of d-rGO-B at different concentrations and of composite films when in direct contact with bacteria (Table S2 and Table S3). Importantly, the d-rGO-B ME-AM scaffolds also demonstrated higher antibacterial activity with increasing d-rGO-B concentration within the scaffolds (Figure 6A). Overall, scaffolds showed to be more potent against *P. aeruginosa* (Gram -) than *S.epidermidis* (Gram +). Interestingly, antibacterial effects were only evident when bacteria were placed in direct contact with the scaffolds, and no biocidal effect of the supernatant, in which scaffolds were incubated up to 3 days (Figure S7), was observed. This suggests that reactive oxygen species (ROS)-dependent oxidative stress was not the dominant antibacterial mechanism, but instead the mechanism was membrane stress induced by sharp edges or corners of d-rGO-B exposed to the surface of the scaffolds, which act as nano-knives or nanoneedles and disrupt bacteria membranes. [19, 58, 63] Moreover, this explains the increasing antimicrobial activity with increasing d-rGO-B concentration, and therefore more d-rGO-B exposed to the filaments surface. In addition, rGO produces lower oxidative stress compared to GO, because of the lower amount of oxygen functional groups [64] Due to the conductivity of rGO, charge-transfer oxidative stress can also be considered a main antimicrobial mechanism when bacteria get in contact with the surface of the scaffolds. Here, rGO can act as a conductive bridge over the bacteria lipid bilayer to mediate electron transfer from bacterial intracellular components to the external environment, interrupting the bacteria membrane respiratory chain, and leading to bacteria death. [63, 64]

**Figure 6.**
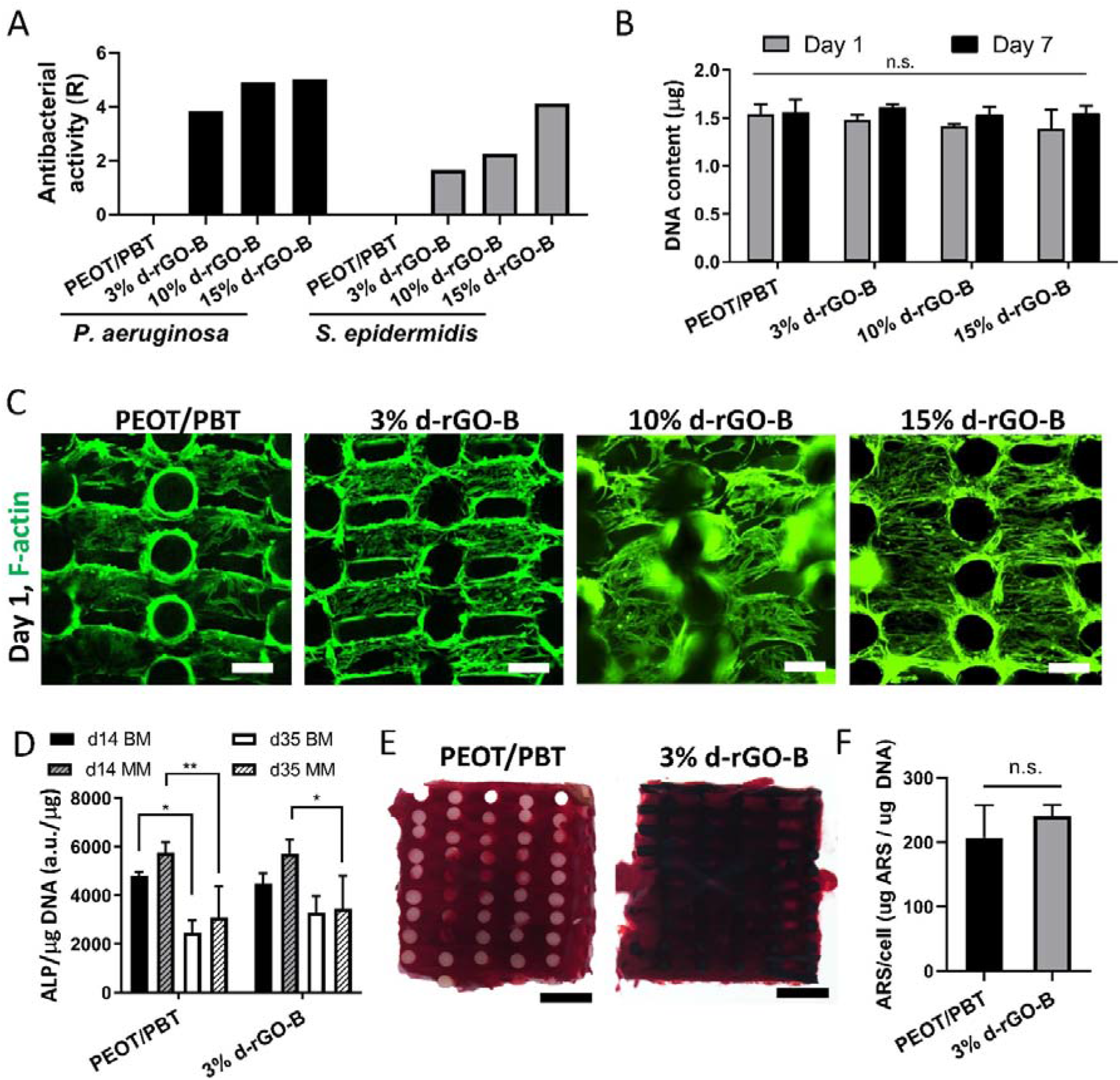
Cell-material interactions in d-rGO-B scaffolds: effect of d-rGO-B concentration. (A) Antibacterial activity of scaffolds against P. aeruginosa and S. epidermidis. (B) DNA content on scaffolds seeded with hMSCs after 1 and 7 days of culture. (C) Fluorescent microscopy images of hMSCs attachment (F-actin, green) to composite scaffolds cross section. Scale bars 250 μm. (D) ALP activity of hMSCs seeded on PEOT/PBT and 3% d-rGO-B scaffolds after 14 and 35 days cultured in BM or MM. Data presented as average ± s.d. and statistical significance performed using two-way ANOVA with Tukey’s multiple comparison test (* p < 0.05, ** p < 0.01). (E) Stereomicroscopy images of PEOT/PBT and 3% d-rGO-B scaffold cross sections stained with alizarin red S after 35 days of culture in MM. Scale bars 1 mm. (F) Quantification of the alizarin red S staining extracted from scaffolds in (E), normalized to cell number. Data presented as average ± s.d. and statistical significance performed using T-test (n.s. p > 0.05).

#### Cell seeding on scaffolds and osteogenic differentiation

In spite of scaffolds displaying antibacterial activity, hMSCs viability was not compromised when seeded on the d-rGO-B composite scaffolds, at any d-rGO-B concentration. As presented in Figure 6C, cells populated the scaffolds filaments from the beginning of the culture, and showed viable cells characteristics, such as spread or elongated morphology. DNA analysis further confirmed the biocompatibility of the scaffolds up to 15% d-rGO-B, as cell number did not decrease after 7 days of culture, but was maintained rather constant (Figure 6B). While various reports have addressed the cyto- or genotoxicity of graphene and GO in contact with different cell lines, [65–67] only a few have tried to understand the biocompatibility of rGO, which greatly differs from the other derivatives in terms crucial factors for biointeractions, such as size (number of layers, available surface area), functional groups (reduction degree) and protein interactions. It has been demonstrated that the cytotoxic effect of GO and partially reduced rGO can be attenuated by proteins in serum binding to their surface, which can reduce the GO/rGO ability to penetrate or physically damage the cell membrane, both in *in vitro* cell cultures and *in vivo.* [68] In addition, rGO has been found to be, in general, more biocompatible than GO and to provide a reduced inflammatory response upon *in vivo* implantation, [69] but still its dose- and size-dependent cytotoxicity towards hMSCs has been shown. [70] Due to the d-rGO-B encapsulated within the polymer matrix and the polymer slow degradation rate such a cytotoxic effect was not evidenced during the culture period addressed here (35 days). [71, 72] Future studies on scaffolds degradation kinetics and d-rGO-B release rate will be essential, as polymer degradation will eventually occur over time in an *in vivo* situation, releasing potentially toxic amounts of d-rGO-B.

Using BM and MM culture medium, preliminary osteogenic differentiation studies were performed with hMSCs seeded on 3% d-rGO-B scaffolds, a low concentration graphene containing scaffold that has been commonly studied in previous reports. [55, 73–75] HMSCs were able to proliferate both in PEOT/PBT and 3% d-rGO-B scaffolds over the culture period (35 days) especially in MM, likely due to more ECM formation within the scaffolds pores acting as cell growth support, compared to BM (Figure S8). As an early osteogenic marker, Figure 6D demonstrates the ALP activity of hMSCs scaffolds. For both bare PEOT/PBT and 3 % d-rGO-B containing scaffolds, a peak in ALP activity was shown at day 14, with a significant decrease on day 35, both in BM and MM, which is in agreement with the osteogenic differentiation progression. [76] However, no statistical differences on ALP activity were found among the d-rGO-B containing scaffolds and the PEOT/PBT control. ECM mineralization was examined at day 35 of culture in BM and MM with ARS, as a late osteogenic marker. Interestingly, both PEOT/PBT and 3% d-rGO-B scaffolds revealed calcium deposition on the ECM matrix produced by cells inside the pores of scaffolds cultured in MM, as depicted by the ARS in Figure 6E. In spite of a slight increase in the calcium deposition on 3% d-rGO-B scaffolds, no statistical differences were observed among these and the PEOT/PBT scaffolds (Figure 6F). These results imply that 3% d-rGO-B scaffolds support osteogenic differentiation, observed by both early and late stage markers. Yet, these scaffolds seem to not enhance differentiation compared to bare polymeric scaffolds. While this is in disagreement with some previous reports, other studies only point out the enhanced cell adhesion and proliferation ability on composite scaffolds with such low graphene derivatives concentrations (up to 1 wt%). [16, 29, 73] As previously mentioned, this might very well be correlated to differences in rGO bulk density affecting the final volume fraction of rGO within the scaffolds. It is plausible that the low d-rGO-B concentration used here (3 wt%) did not allow for sufficient exposure of d-rGO-B to the scaffolds filament surface, preventing it to act as a pre-concentration platform for osteogenic inducers (e.g. dexamethasone and β-glycerolphosphate), which is believed to be the origin of the enhanced stem cell differentiation exerted by graphene derivatives. [77–79] In this regard, we hypothesize that the increased hydrophilicity and protein adsorption on 10% and 15% d-rGO-B scaffolds, together with their surface microroughness, would induce accelerated osteogenic differentiation, compared to 3% d-rGO-B and PEOT/PBT scaffolds. Nevertheless, future studies will be dedicated to validate this hypothesis.

## 4. Conclusions

Due to its very low bulk density, rGO has to be densified prior to melt compounding with a polymer. The aim of this study was to understand the effect of rGO densification parameters (densification solvent and rGO bulk density after densification) on rGO compaction degree and on the resulting melt compounded composites physicochemical properties and printability via ME-AM. The effect of rGO concentration on the materials’ physicochemical properties and cell-material interactions in 2D and 3D was also investigated. It was observed that high d-rGO bulk density (90 g/L) correlated with higher d-rGO compaction, which was translated into smaller d-rGO volume fraction for a given d-rGO concentration within the polymer composite, and printability up to 15 wt% d-rGO. On the other hand, medium bulk density d-rGO (50 g/L) occupied a greater volume within the melt compounded composites, which presented some challenges upon ME-AM at high d-rGO contents (10 and 15 wt%), in terms of loss of filament fidelity, poor layer bonding, or even lack of printability. At a given bulk density (50 g/L), when comparing acetone and water as densification solvent, it was observed that densification in water led to a more compacted d-rGO than d-rGO densified in acetone, and d-rGO was poorly dispersed within the polymer matrix. This led to bigger aggregates formation within the composites, whose connections were able to create conductive pathways within the composites, making these materials the most conductive among all d-rGO types. Composites prepared medium bulk density d-rGO-B densified in acetone were chosen for further characterization, due to a balance of printability and electrical properties. It was observed that increasing d-rGO content led to increasing material hydrophilicity and protein adsorption, as well as to increasing surface roughness due to higher rGO exposure to the surface. Scaffolds prepared with 3, 10 and 15% d-rGO were found to possess increasing antibacterial properties with increased d-rGO content, without affecting hMSCs viability. Notably, 3% d-rGO scaffolds were able to support hMSCs proliferation and osteogenic differentiation. Overall, this study demonstrated that rGO compaction degree and concentration greatly affects composites printability and scaffolds physicochemical and electrical properties. In this regard, a careful selection of the rGO densification parameters has to be made in order to ensure the most adequate properties of the final scaffolds required for bone tissue engineering applications.

## Acknowledgements

We are grateful to the FAST project funded under the H2020-NMP-PILOTS-2015 scheme (GA n. 685825) for financial support. Some of the materials used in this work were provided by the Texas A&M Health Science Center College of Medicine Institute for Regenerative Medicine at Scott & White through a grant from NCRR of the NIH (Grant #P40RR017447).

## References

[1] Z.M. Wright, A.M. Arnold, B.D. Holt, K.E. Eckhart, S.A. Sydlik, Functional Graphenic Materials, Graphene Oxide, and Graphene as Scaffolds for Bone Regeneration, Regenerative Engineering and Translational Medicine 5(2) (2019) 190–209.

[2] S.R. Shin, Y.-C. Li, H.L. Jang, P. Khoshakhlagh, M. Akbari, A. Nasajpour, Y.S. Zhang, A. Tamayol, A. Khademhosseini, Graphene-based materials for tissue engineering, Advanced Drug Delivery Reviews.

[3] M. Gu, Y. Liu, T. Chen, F. Du, X. Zhao, C. Xiong, Y. Zhou, Is Graphene a Promising Nano-Material for Promoting Surface Modification of Implants or Scaffold Materials in Bone Tissue Engineering?, Tissue Engineering. Part B, Reviews 20(5) (2014) 477–491.

[4] C. Lee, X. Wei, J.W. Kysar, J. Hone, Measurement of the elastic properties and intrinsic strength of monolayer graphene, science 321(5887) (2008) 385–388.

[5] X. Du, I. Skachko, A. Barker, E.Y. Andrei, Approaching ballistic transport in suspended graphene, Nature Nanotechnology 3(8) (2008) 491–495.

[6] K.P. Loh, Q. Bao, P.K. Ang, J. Yang, The chemistry of graphene, Journal of Materials Chemistry 20(12) (2010) 2277–2289.

[7] Y. Wang, X. Chen, Y. Zhong, F. Zhu, K.P. Loh, Large area, continuous, few-layered graphene as anodes in organic photovoltaic devices, Applied Physics Letters 95(6) (2009) 063302.

[8] K.S. Novoselov, A.K. Geim, S.V. Morozov, D. Jiang, Y. Zhang, S.V. Dubonos, I.V. Grigorieva, A.A. Firsov, Electric field effect in atomically thin carbon films, science 306(5696) (2004) 666–669.

[9] D.R. Dreyer, S. Park, C.W. Bielawski, R.S. Ruoff, The chemistry of graphene oxide, Chemical Society Reviews 39(1) (2010) 228–240.

[10] T. Kuila, S. Bose, A.K. Mishra, P. Khanra, N.H. Kim, J.H. Lee, Chemical functionalization of graphene and its applications, Progress in Materials Science 57(7) (2012) 1061–1105.

[11] S. Pei, H.-M. Cheng, The reduction of graphene oxide, Carbon 50(9) (2012) 3210–3228.

[12] F. Rostami, E. Tamjid, M. Behmanesh, Drug-eluting PCL/graphene oxide nanocomposite scaffolds for enhanced osteogenic differentiation of mesenchymal stem cells, Materials Science and Engineering: C 115 (2020) 111102.

[13] T. Zhou, G. Li, S. Lin, T. Tian, Q. Ma, Q. Zhang, S. Shi, C. Xue, W. Ma, X. Cai, Y. Lin, Electrospun Poly(3-hydroxybutyrate-co-4-hydroxybutyrate)/Graphene Oxide Scaffold: Enhanced Properties and Promoted in Vivo Bone Repair in Rats, ACS Applied Materials & Interfaces 9(49) (2017) 42589–42600.

[14] C. Cha, S.R. Shin, X. Gao, N. Annabi, M.R. Dokmeci, X.S. Tang, A. Khademhosseini, Controlling mechanical properties of cell-laden hydrogels by covalent incorporation of graphene oxide, Small 10(3) (2014) 514–523.

[15] J. Li, X. Liu, J.M. Crook, G.G. Wallace, 3D Printing of Cytocompatible Graphene/Alginate Scaffolds for Mimetic Tissue Constructs, Frontiers in Bioengineering and Biotechnology 8(824) (2020).

[16] W. Wang, G. Caetano, W.S. Ambler, J.J. Blaker, M.A. Frade, P. Mandal, C. Diver, P. Bartolo, Enhancing the Hydrophilicity and Cell Attachment of 3D Printed PCL/Graphene Scaffolds for Bone Tissue Engineering, Materials (Basel, Switzerland) 9(12) (2016).

[17] H. Belaid, S. Nagarajan, C. Teyssier, C. Barou, J. Barés, S. Balme, H. Garay, V. Huon, D. Cornu, V. Cavaillès, M. Bechelany, Development of new biocompatible 3D printed graphene oxide-based scaffolds, Materials Science and Engineering: C 110 (2020) 110595.

[18] A.E. Jakus, E.B. Secor, A.L. Rutz, S.W. Jordan, M.C. Hersam, R.N. Shah, Three-dimensional printing of high-content graphene scaffolds for electronic and biomedical applications, ACS nano 9(4) (2015) 4636–4648.

[19] C. Angulo-Pineda, K. Srirussamee, P. Palma, V.M. Fuenzalida, S.H. Cartmell, H. Palza, Electroactive 3D Printed Scaffolds Based on Percolated Composites of Polycaprolactone with Thermally Reduced Graphene Oxide for Antibacterial and Tissue Engineering Applications, Nanomaterials 10(3) (2020) 428.

[20] J.R. Potts, D.R. Dreyer, C.W. Bielawski, R.S. Ruoff, Graphene-based polymer nanocomposites, Polymer 52(1) (2011) 5–25.

[21] A.L. Higginbotham, J.R. Lomeda, A.B. Morgan, J.M. Tour, Graphite oxide flame-retardant polymer nanocomposites, ACS Appl Mater Interfaces 1(10) (2009) 2256–61.

[22] F.d.C. Fim, J.M. Guterres, N.R. Basso, G.B. Galland, Polyethylene/graphite nanocomposites obtained by in situ polymerization, Journal of Polymer Science Part A: Polymer Chemistry 48(3) (2010) 692–698.

[23] D.R. Paul, L.M. Robeson, Polymer nanotechnology: Nanocomposites, Polymer 49(15) (2008) 3187–3204.

[24] H. Kim, S. Kobayashi, M.A. AbdurRahim, M.J. Zhang, A. Khusainova, M.A. Hillmyer, A.A. Abdala, C.W. Macosko, Graphene/polyethylene nanocomposites: Effect of polyethylene functionalization and blending methods, Polymer 52(8) (2011) 1837–1846.

[25] Q. Chen, J.D. Mangadlao, J. Wallat, A. De Leon, J.K. Pokorski, R.C. Advincula, 3D Printing Biocompatible Polyurethane/Poly(lactic acid)/Graphene Oxide Nanocomposites: Anisotropic Properties, ACS Applied Materials & Interfaces 9(4) (2017) 4015–4023.

[26] M.J. McAllister, J.-L. Li, D.H. Adamson, H.C. Schniepp, A.A. Abdala, J. Liu, M. Herrera-Alonso, D.L. Milius, R. Car, R.K. Prud’homme, I.A. Aksay, Single Sheet Functionalized Graphene by Oxidation and Thermal Expansion of Graphite, Chemistry of Materials 19(18) (2007) 4396–4404.

[27] K. Kalaitzidou, H. Fukushima, L.T. Drzal, A new compounding method for exfoliated graphite–polypropylene nanocomposites with enhanced flexural properties and lower percolation threshold, Composites Science and Technology 67(10) (2007) 2045–2051.

[28] P. Steurer, R. Wissert, R. Thomann, R. Mülhaupt, Functionalized Graphenes and Thermoplastic Nanocomposites Based upon Expanded Graphite Oxide, Macromolecular Rapid Communications 30(4-5) (2009) 316–327.

[29] W. Wang, G.F. Caetano, W.-H. Chiang, A.L. Braz, J.J. Blaker, M.A.C. Frade, P.J.D.S. Bartolo, Morphological, mechanical and biological assessment of PCL/pristine graphene scaffolds for bone regeneration, International Journal of Bioprinting 2(2) (2016).

[30] W. Wang, J.R.P. Junior, P.R.L. Nalesso, D. Musson, J. Cornish, F. Mendonça, G.F. Caetano, P. Bártolo, Engineered 3D printed poly(J-caprolactone)/graphene scaffolds for bone tissue engineering, Materials Science and Engineering: C 100 (2019) 759–770.

[31] Y. Hou, W. Wang, P. Bártolo, Novel Poly (J-caprolactone)/Graphene Scaffolds for Bone Cancer Treatment and Bone Regeneration, 3D Printing and Additive Manufacturing (2020).

[32] R. Sinha, M. Cámara-Torres, P. Scopece, E.V. Falzacappa, A. Patelli, L. Moroni, C. Mota, A hybrid additive manufacturing platform to create bulk and surface composition gradients on scaffolds for tissue regeneration, bioRxiv (2020) 2020.06.23.165605.

[33] M. Cámara-Torres, R. Sinha, C. Mota, L. Moroni, Improving cell distribution on 3D additive manufactured scaffolds through engineered seeding media density and viscosity, Acta Biomaterialia 101 (2020) 183–195.

[34] A.V. Rane, K. Kanny, V.K. Abitha, S. Thomas, Chapter 5 - Methods for Synthesis of Nanoparticles and Fabrication of Nanocomposites, in: S. Mohan Bhagyaraj, O.S. Oluwafemi, N. Kalarikkal, S. Thomas (Eds.), Synthesis of Inorganic Nanomaterials, Woodhead Publishing2018, pp. 121–139.

[35] H. Kim, A.A. Abdala, C.W. Macosko, Graphene/Polymer Nanocomposites, Macromolecules 43(16) (2010) 6515–6530.

[36] S. Araby, I. Zaman, Q. Meng, N. Kawashima, A. Michelmore, H.-C. Kuan, P. Majewski, J. Ma, L. Zhang, Melt compounding with graphene to develop functional, high-performance elastomers, Nanotechnology 24(16) (2013) 165601.

[37] X. Zhang, D.V. Raj, X. Zhou, Z. Liu, Solvent evaporation induced graphene powder with high volumetric capacitance and outstanding rate capability for supercapacitors, Journal of Power Sources 382 (2018) 95–100.

[38] H.L. Poh, F. Šaněk, A. Ambrosi, G. Zhao, Z. Sofer, M. Pumera, Graphenes prepared by Staudenmaier, Hofmann and Hummers methods with consequent thermal exfoliation exhibit very different electrochemical properties, Nanoscale 4(11) (2012) 3515–3522.

[39] A. Dimiev, D.V. Kosynkin, L.B. Alemany, P. Chaguine, J.M. Tour, Pristine Graphite Oxide, Journal of the American Chemical Society 134(5) (2012) 2815–2822.

[40] J. Guerrero-Contreras, F. Caballero-Briones, Graphene oxide powders with different oxidation degree, prepared by synthesis variations of the Hummers method, Materials Chemistry and Physics 153 (2015) 209–220.

[41] C.H.A. Wong, Z. Sofer, M. Kubešová, J. Kučera, S. Matějková, M. Pumera, Synthetic routes contaminate graphene materials with a whole spectrum of unanticipated metallic elements, Proceedings of the National Academy of Sciences 111(38) (2014) 13774.

[42] V.B. Mohan, R. Brown, K. Jayaraman, D. Bhattacharyya, Characterisation of reduced graphene oxide: Effects of reduction variables on electrical conductivity, Materials Science and Engineering: B 193 (2015) 49–60.

[43] N. Díez, A. Śliwak, S. Gryglewicz, B. Grzyb, G. Gryglewicz, Enhanced reduction of graphene oxide by high-pressure hydrothermal treatment, Rsc Advances 5(100) (2015) 81831–81837.

[44] S. Joshi, R. Siddiqui, P. Sharma, R. Kumar, G. Verma, A. Saini, Green synthesis of peptide functionalized reduced graphene oxide (rGO) nano bioconjugate with enhanced antibacterial activity, Scientific Reports 10(1) (2020) 9441.

[45] M. Simón, A. Benítez, A. Caballero, J. Morales, O. Vargas, Untreated Natural Graphite as a Graphene Source for High-Performance Li-Ion Batteries, Batteries 4(1) (2018).

[46] K.-H. Liao, Y.T. Park, A. Abdala, C. Macosko, Aqueous reduced graphene/thermoplastic polyurethane nanocomposites, Polymer 54(17) (2013) 4555–4559.

[47] X. Zhao, Q. Zhang, D. Chen, P. Lu, Enhanced Mechanical Properties of Graphene-Based Poly(vinyl alcohol) Composites, Macromolecules 43(5) (2010) 2357–2363.

[48] C. Shuai, P. Feng, C. Gao, X. Shuai, T. Xiao, S. Peng, Graphene oxide reinforced poly(vinyl alcohol): nanocomposite scaffolds for tissue engineering applications, RSC Advances 5(32) (2015) 25416–25423.

[49] W.E. Mahmoud, Morphology and physical properties of poly(ethylene oxide) loaded graphene nanocomposites prepared by two different techniques, European Polymer Journal 47(8) (2011) 1534–1540.

[50] B. Shen, W. Zhai, M. Tao, D. Lu, W. Zheng, Enhanced interfacial interaction between polycarbonate and thermally reduced graphene induced by melt blending, Composites Science and Technology 86 (2013) 109–116.

[51] B. Shen, W. Zhai, C. Chen, D. Lu, J. Wang, W. Zheng, Melt Blending In situ Enhances the Interaction between Polystyrene and Graphene through π–π Stacking, ACS Applied Materials & Interfaces 3(8) (2011) 3103–3109.

[52] A.J. Marsden, D.G. Papageorgiou, C. Vallés, A. Liscio, V. Palermo, M.A. Bissett, R.J. Young, I.A. Kinloch, Electrical percolation in graphene–polymer composites, 2D Materials 5(3) (2018) 032003.

[53] K. Huang, J. Yang, S. Dong, Q. Feng, X. Zhang, Y. Ding, J. Hu, Anisotropy of graphene scaffolds assembled by three-dimensional printing, Carbon 130 (2018) 1–10.

[54] R. Sinha, A. Sanchez, M. Camara-Torres, I.C. Uriszar-Aldaca, A.R. Calore, J. Harings, A. Gambardella, L. Ciccarelli, V. Vanzanella, M. Sisani, M. Scatto, R. Wendelbo, S. Perez, S. Villanueva, A. Matanza, A. Patelli, N. Grizzuti, C. Mota, L. Moroni, Additive manufactured scaffolds for bone tissue engineering: physical characterization of thermoplastic composites with functional fillers, bioRxiv (2021) 2021.03.23.436548.

[55] W. Wang, B. Huang, J.J. Byun, P. Bártolo, Assessment of PCL/carbon material scaffolds for bone regeneration, Journal of the mechanical behavior of biomedical materials 93 (2019) 52–60.

[56] S. Pei, F. Ai, S. Qu, Fabrication and biocompatibility of reduced graphene oxide/poly(vinylidene fluoride) composite membranes, RSC Advances 5(121) (2015) 99841–99847.

[57] P. Arriagada, H. Palza, P. Palma, M. Flores, P. Caviedes, Poly(lactic acid) composites based on graphene oxide particles with antibacterial behavior enhanced by electrical stimulus and biocompatibility, Journal of Biomedical Materials Research Part A 106(4) (2018) 1051–1060.

[58] S. Kumar, S. Raj, E. Kolanthai, A.K. Sood, S. Sampath, K. Chatterjee, Chemical Functionalization of Graphene To Augment Stem Cell Osteogenesis and Inhibit Biofilm Formation on Polymer Composites for Orthopedic Applications, ACS Applied Materials & Interfaces 7(5) (2015) 3237–3252.

[59] S. Kumar, S.H. Parekh, Linking graphene-based material physicochemical properties with molecular adsorption, structure and cell fate, Communications Chemistry 3(1) (2020) 8.

[60] X. Shi, H. Chang, S. Chen, C. Lai, A. Khademhosseini, H. Wu, Regulating Cellular Behavior on Few-Layer Reduced Graphene Oxide Films with Well-Controlled Reduction States, Advanced Functional Materials 22(4) (2012) 751–759.

[61] J.Y. Lim, J.C. Hansen, C.A. Siedlecki, J. Runt, H.J. Donahue, Human foetal osteoblastic cell response to polymer-demixed nanotopographic interfaces, Journal of The Royal Society Interface 2(2) (2005) 97–108.

[62] A. Patelli, F. Mussano, P. Brun, T. Genova, E. Ambrosi, N. Michieli, G. Mattei, P. Scopece, L. Moroni, Nanoroughness, Surface Chemistry, and Drug Delivery Control by Atmospheric Plasma Jet on Implantable Devices, ACS Applied Materials & Interfaces 10(46) (2018) 39512–39523.

[63] Z. Jia, Y. Shi, P. Xiong, W. Zhou, Y. Cheng, Y. Zheng, T. Xi, S. Wei, From Solution to Biointerface: Graphene Self-Assemblies of Varying Lateral Sizes and Surface Properties for Biofilm Control and Osteodifferentiation, ACS Applied Materials & Interfaces 8(27) (2016) 17151–17165.

[64] X. Zou, L. Zhang, Z. Wang, Y. Luo, Mechanisms of the Antimicrobial Activities of Graphene Materials, Journal of the American Chemical Society 138(7) (2016) 2064–2077.

[65] K. Wang, J. Ruan, H. Song, J. Zhang, Y. Wo, S. Guo, D. Cui, Biocompatibility of Graphene Oxide, Nanoscale Res Lett 6(1) (2010) 8.

[66] B.D. Holt, A.M. Arnold, S.A. Sydlik, In It for the Long Haul: The Cytocompatibility of Aged Graphene Oxide and Its Degradation Products, Advanced Healthcare Materials 5(23) (2016) 3056–3066.

[67] S.P. Mukherjee, A.R. Gliga, B. Lazzaretto, B. Brandner, M. Fielden, C. Vogt, L. Newman, A.F. Rodrigues, W. Shao, P.M. Fournier, M.S. Toprak, A. Star, K. Kostarelos, K. Bhattacharya, B. Fadeel, Graphene oxide is degraded by neutrophils and the degradation products are non-genotoxic, Nanoscale 10(3) (2018) 1180–1188.

[68] W. Hu, C. Peng, M. Lv, X. Li, Y. Zhang, N. Chen, C. Fan, Q. Huang, Protein Corona-Mediated Mitigation of Cytotoxicity of Graphene Oxide, ACS Nano 5(5) (2011) 3693–3700.

[69] S.A. Sydlik, S. Jhunjhunwala, M.J. Webber, D.G. Anderson, R. Langer, In Vivo Compatibility of Graphene Oxide with Differing Oxidation States, ACS Nano 9(4) (2015) 3866–3874.

[70] O. Akhavan, E. Ghaderi, A. Akhavan, Size-dependent genotoxicity of graphene nanoplatelets in human stem cells, Biomaterials 33(32) (2012) 8017–8025.

[71] A.A. Deschamps, D.W. Grijpma, J. Feijen, Poly (ethylene oxide)/poly (butylene terephthalate) segmented block copolymers: the effect of copolymer composition on physical properties and degradation behavior, Polymer 42(23) (2001) 9335–9345.

[72] A. Deschamps, A.A. van Apeldoorn, H. Hayen, J.D. de Bruijn, U. Karst, D.W. Grijpma, J. Feijen, In vivo and in vitro degradation of poly (ether ester) block copolymers based on poly (ethylene glycol) and poly (butylene terephthalate), Biomaterials 25(2) (2004) 247–258.

[73] S. Kumar, D. Azam, S. Raj, E. Kolanthai, K.S. Vasu, A.K. Sood, K. Chatterjee, 3D scaffold alters cellular response to graphene in a polymer composite for orthopedic applications, Journal of Biomedical Materials Research Part B: Applied Biomaterials 104(4) (2016) 732–749.

[74] J.M. Unagolla, A.C. Jayasuriya, Enhanced cell functions on graphene oxide incorporated 3D printed polycaprolactone scaffolds, Materials Science and Engineering: C 102 (2019) 1–11.

[75] K. Krukiewicz, D. Putzer, N. Stuendl, B. Lohberger, F. Awaja, Enhanced Osteogenic Differentiation of Human Primary Mesenchymal Stem and Progenitor Cultures on Graphene Oxide/Poly(methyl methacrylate) Composite Scaffolds, Materials (Basel, Switzerland) 13(13) (2020) 2991.

[76] T.A. Owen, M. Aronow, V. Shalhoub, L.M. Barone, L. Wilming, M.S. Tassinari, M.B. Kennedy, S. Pockwinse, J.B. Lian, G.S. Stein, Progressive development of the rat osteoblast phenotype in vitro: reciprocal relationships in expression of genes associated with osteoblast proliferation and differentiation during formation of the bone extracellular matrix, Journal of cellular physiology 143(3) (1990) 420–30.

[77] W.C. Lee, C.H.Y.X. Lim, H. Shi, L.A.L. Tang, Y. Wang, C.T. Lim, K.P. Loh, Origin of Enhanced Stem Cell Growth and Differentiation on Graphene and Graphene Oxide, ACS Nano 5(9) (2011) 7334–7341.

[78] Y. Luo, H. Shen, Y. Fang, Y. Cao, J. Huang, M. Zhang, J. Dai, X. Shi, Z. Zhang, Enhanced Proliferation and Osteogenic Differentiation of Mesenchymal Stem Cells on Graphene Oxide-Incorporated Electrospun Poly(lactic-co-glycolic acid) Nanofibrous Mats, ACS Applied Materials & Interfaces 7(11) (2015) 6331–6339.

[79] T.R. Nayak, H. Andersen, V.S. Makam, C. Khaw, S. Bae, X. Xu, P.-L.R. Ee, J.-H. Ahn, B.H. Hong, G. Pastorin, B. Özyilmaz, Graphene for Controlled and Accelerated Osteogenic Differentiation of Human Mesenchymal Stem Cells, ACS Nano 5(6) (2011) 4670–4678.

