## Supplementary Information for "Effect of reduced graphene oxide (rGO) compaction degree and concentration on rGO-polymer composites printability and cell interactions"

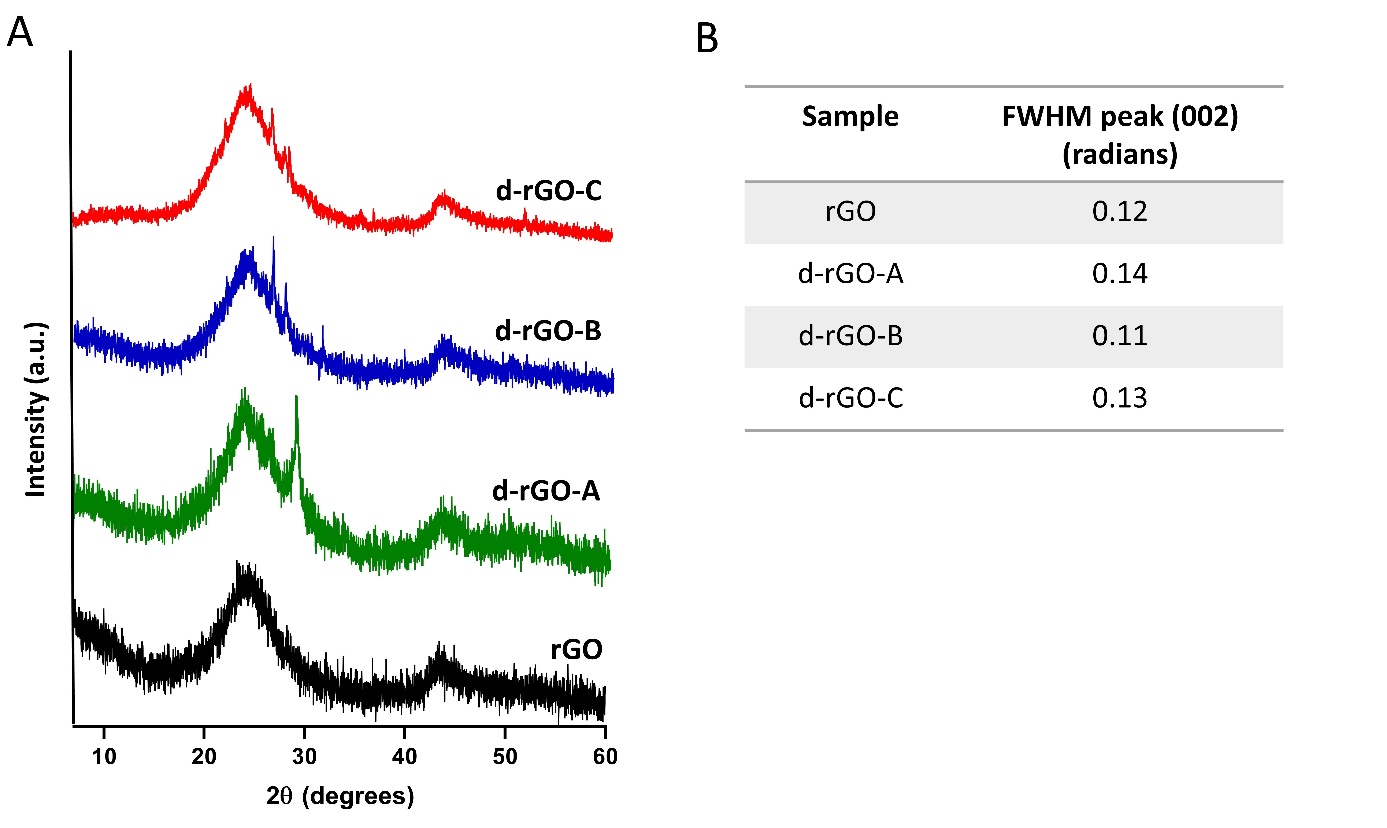

**Figure S1.** (A) XRD diffraction patterns of the three different d-rGO used in this study, each coming from different rGO batches. (B) Full width at half maximum (FWHM) of (002) diffraction peaks.

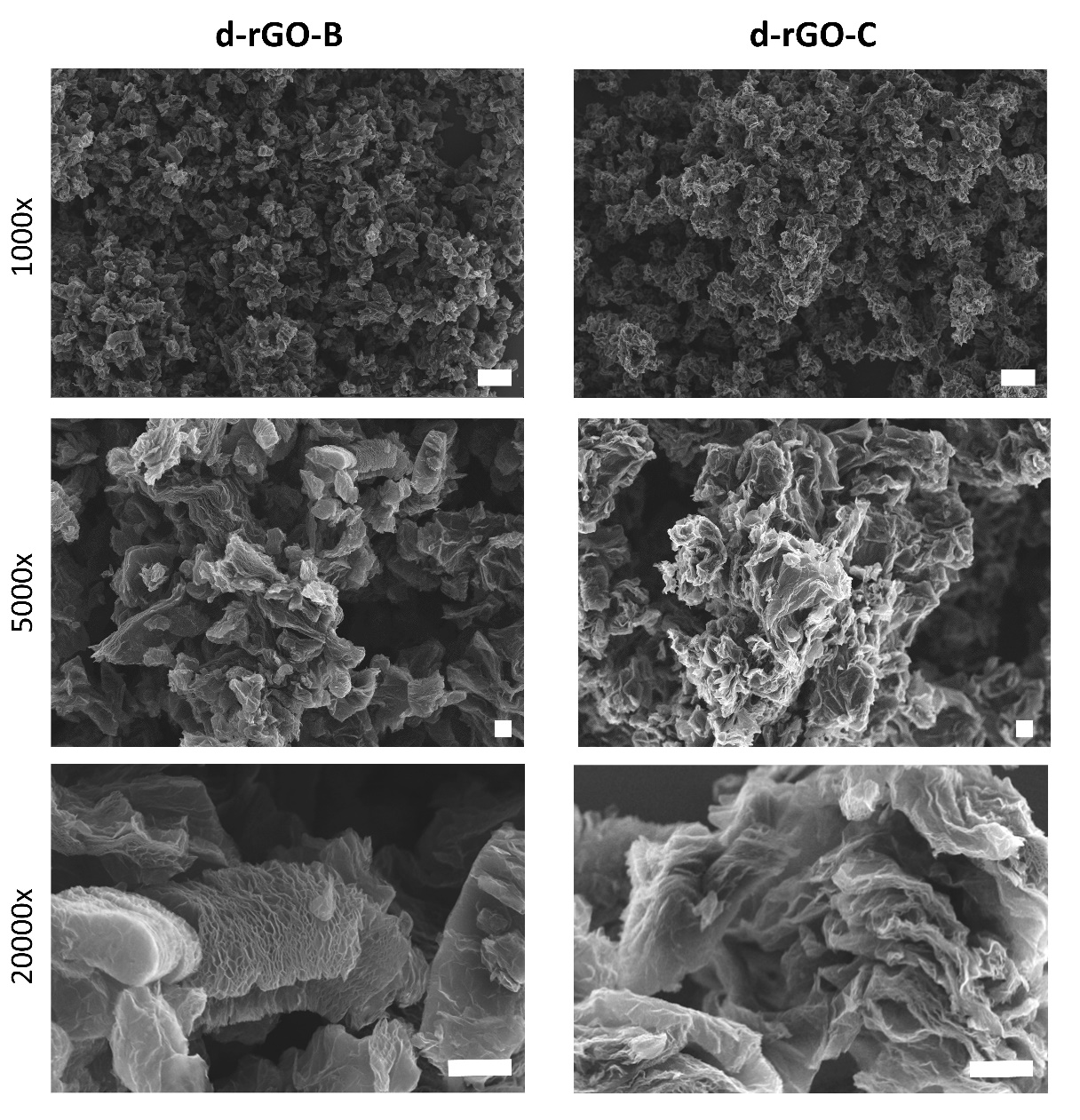

**Figure S2**. Representative SEM images of d-rGO-B and d-rGO-C at different magnifications, displaying different degrees of compaction. Scale bars 20 µm (top row), and 2 µm (middle and bottom row).

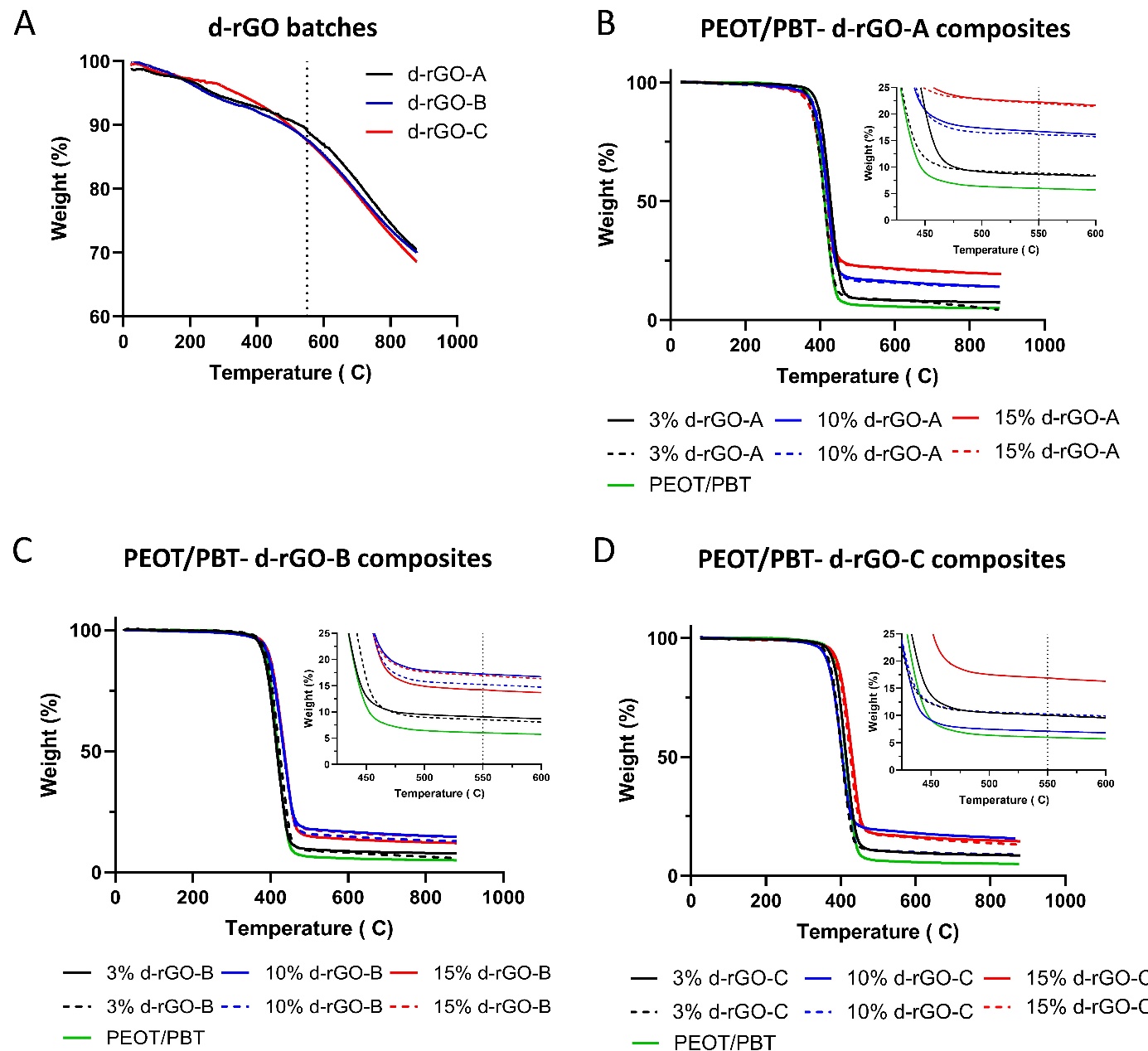

**Figure S3.** TGA curves of (A) d-rGO batches, (B) PEOT/PBT-d-rGO-A composites, (C) PEOT/PBT-d-rGO-B composites and (D) PEOT/PBT-d-rGO-C composites. For each concentration, two curves, measured from two different samples are represented.

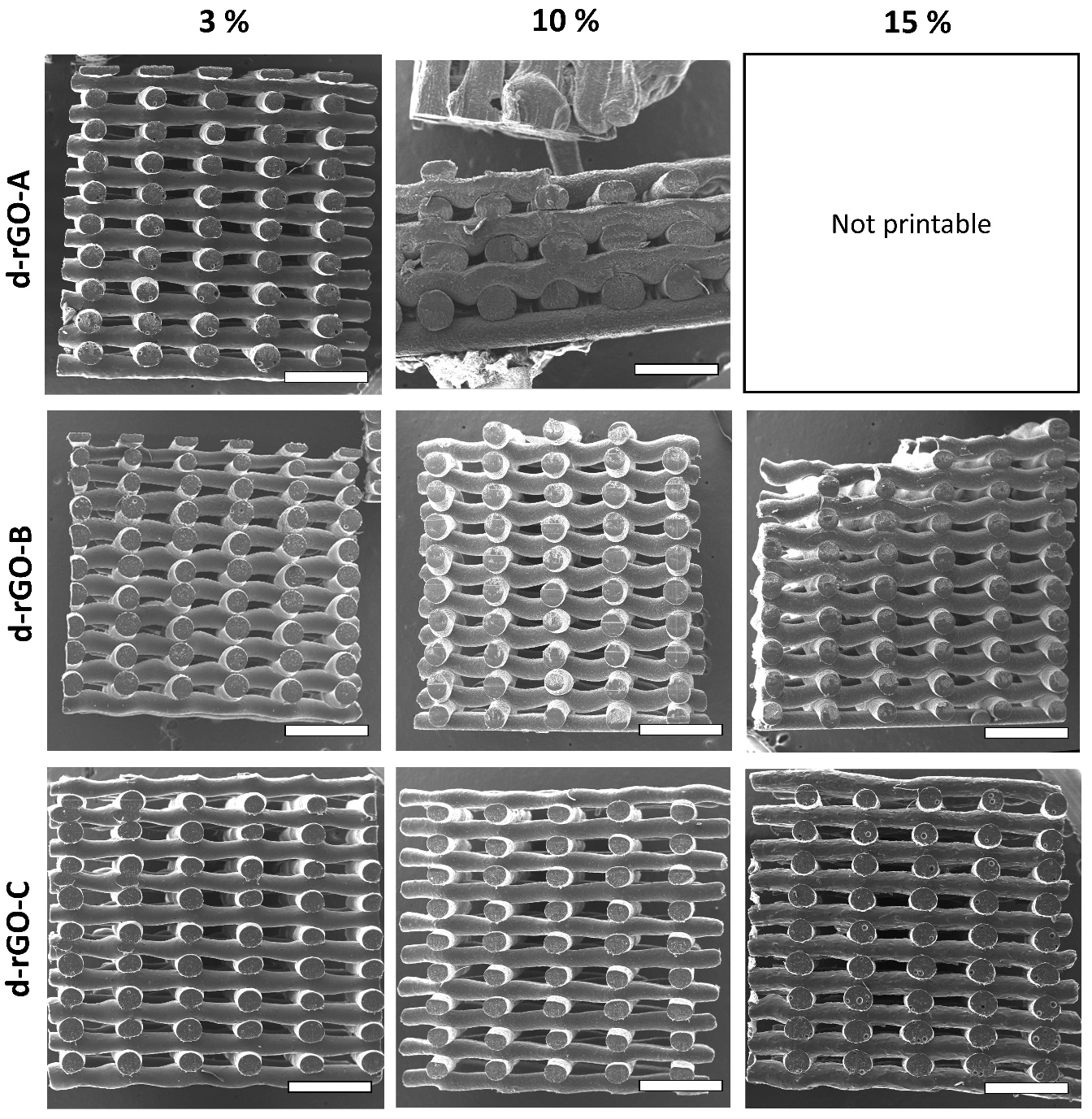

**Figure S4.** SEM micrographs of 3D ME-AM scaffolds cross sections obtained using each of the d-rGO composites, depicting scaffolds morphology and interconnected porosity. Scale bars 1mm.

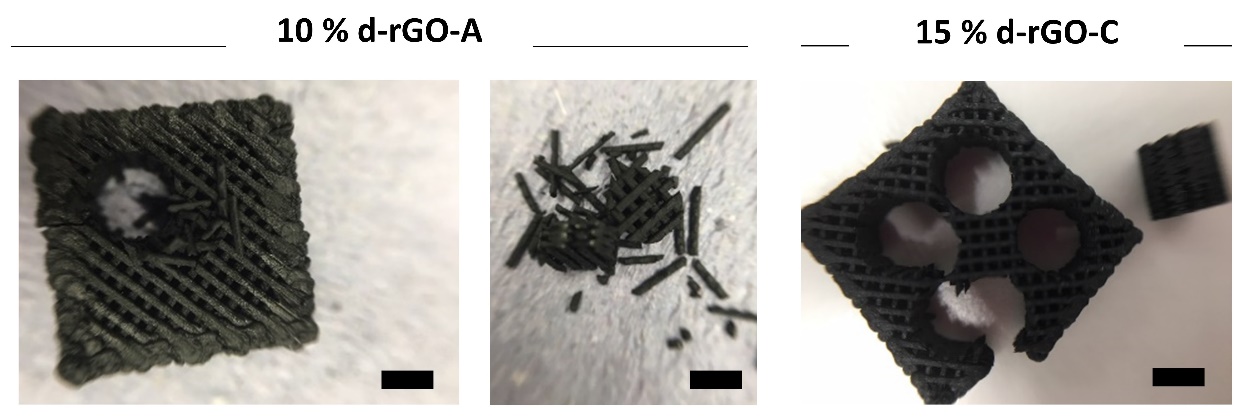

**Figure S5** Demonstration of poor layer bonding of 10% d-rGO-A upon punching of scaffold using a biopsy puncher, compared to other scaffold types, such as the 15% d-rGO-C scaffold. Scale bars 2 mm.

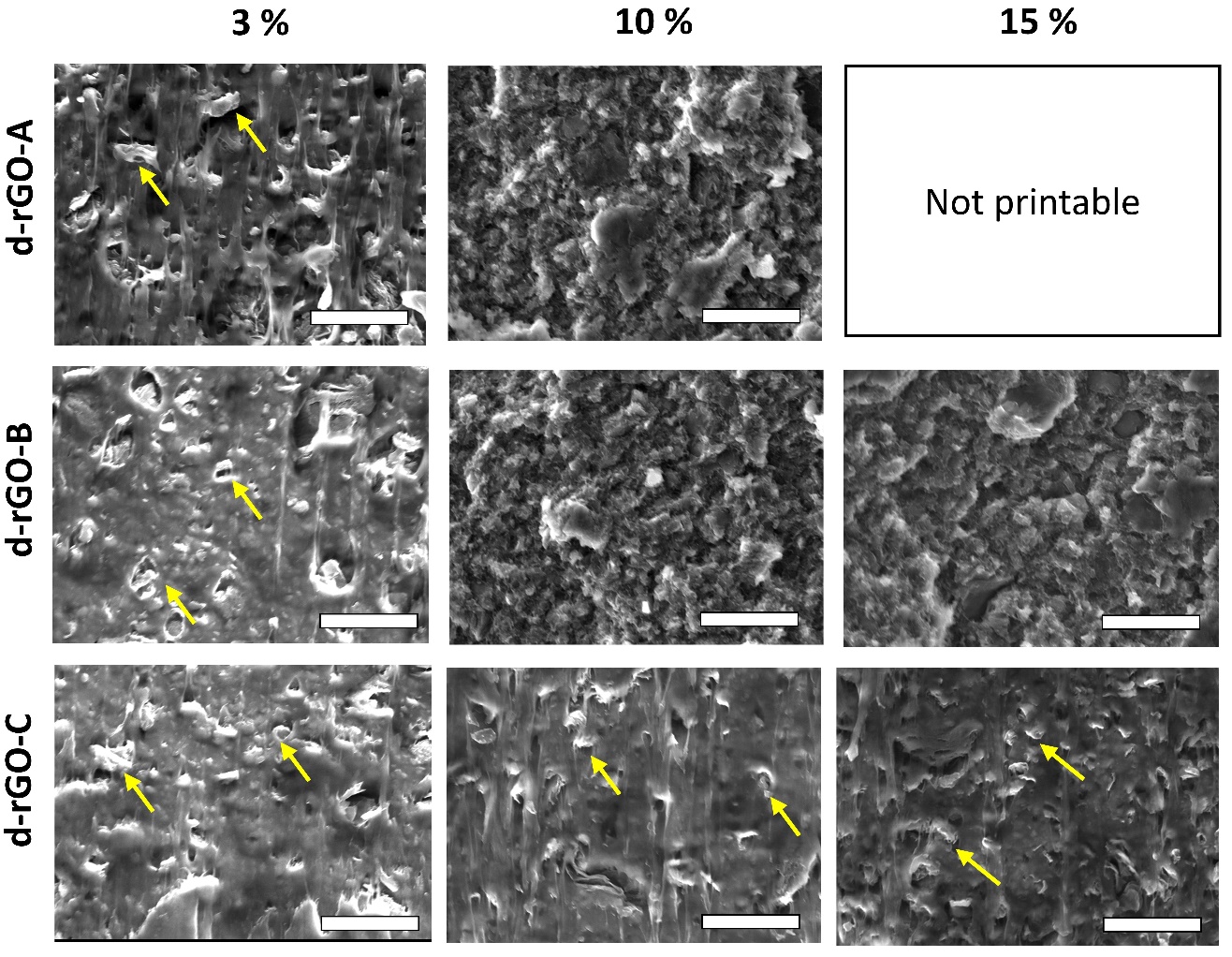

**Figure S6.** SEM micrographs of 3D ME-AM scaffolds filaments cross sections cut with a razor blade. Yellow arrows indicate rGO particles within the polymer matrix. Scale bars 50 µm.

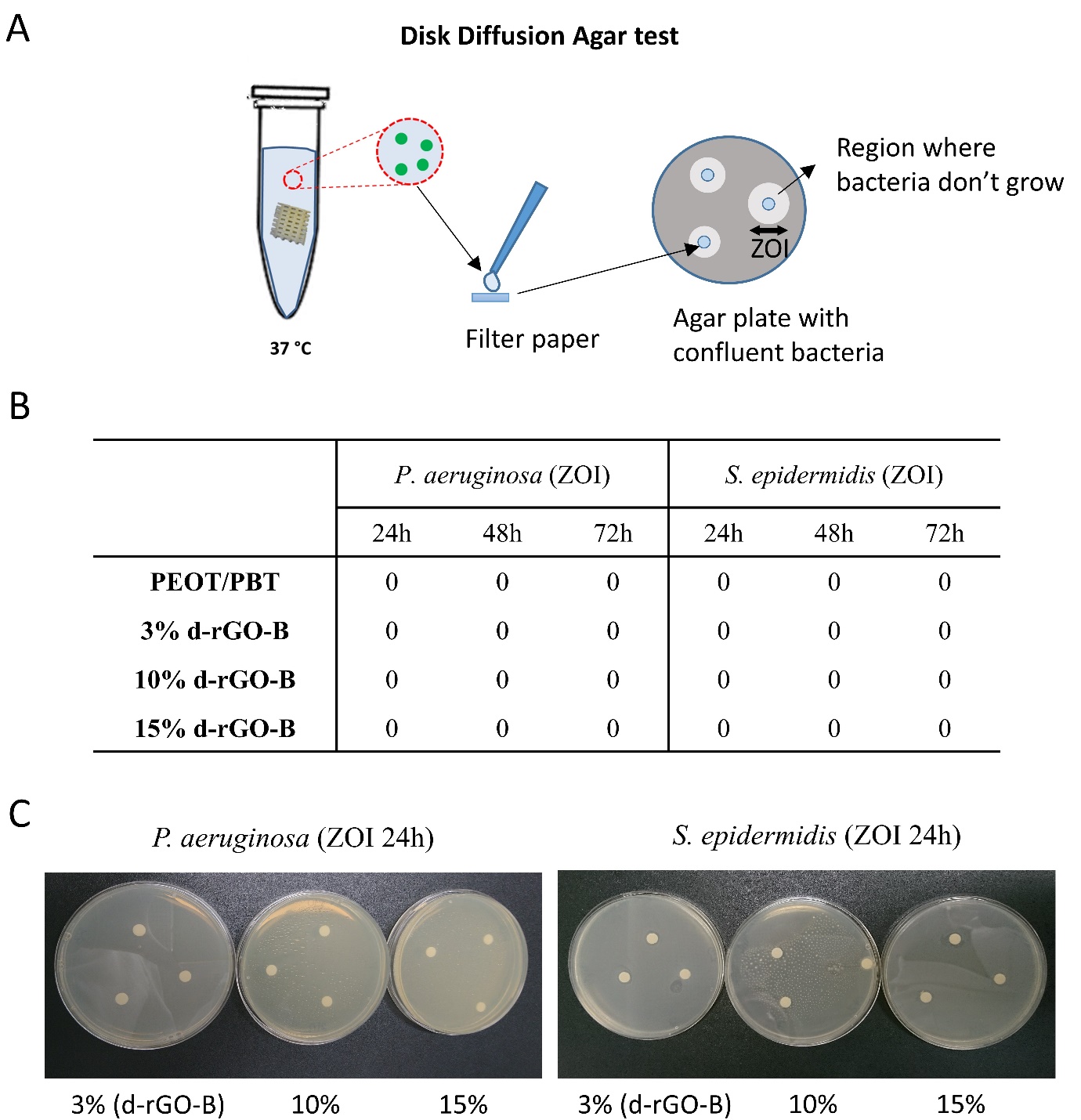

**Figure S7.** (A) Disk diffusion agar test diagram. (B) Antimicrobial activity of d-rGO-B scaffolds against P. aeruginosa and S. epidermidis, measured through the disk diffusion agar test and reported as zone of inhibition (ZOI) values. (C) Images of representative disk diffusion test plates depicting ZOIs around a disk impregnated with an aliquot of the scaffold supernatant after the initial 24h of incubation. Scale bars 5 mm.

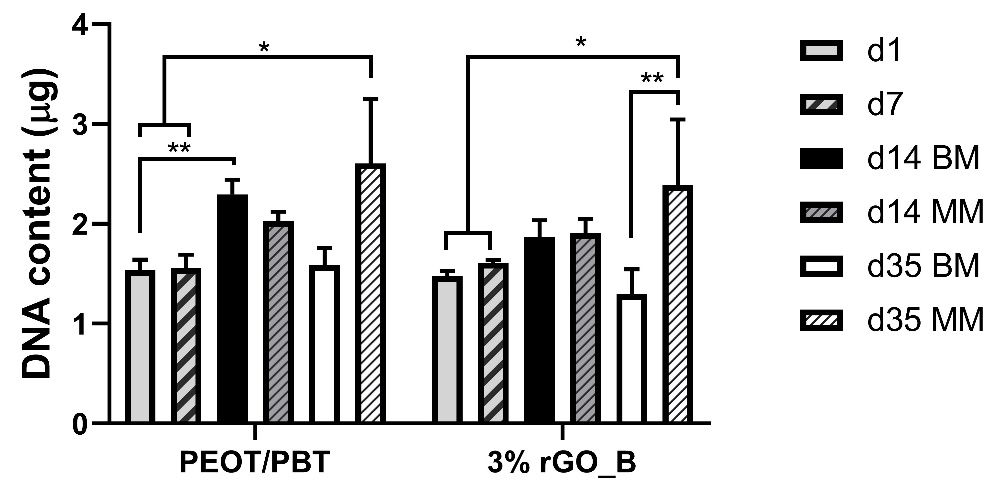

**Figure S8.** DNA content progression on PEOT/PBT and 3% d-rGO-B scaffolds over 35 days of culture in BM or MM.

**Table S1.** Atomic compositions (%) of each of the d-rGO measured by XPS.

|  | C | O | C/O ratio | N | Al | Si | S | Cl | Fe |
| --- | --- | --- | --- | --- | --- | --- | --- | --- | --- |
| **d-rGO-A** | 85.3 | 14.4 | 5.8 | 0.0 | 0.0 | 0.2 | 0.0 | 0.1 | 0.0 |
| **d-rGO-B** | 83 | 16 | 5.2 | 0.2 | 0.0 | 0.6 | 0.2 | 0.2 | 0.0 |
| **d-rGO-C** | 82 | 16.5 | 5.0 | 0.2 | 0.2 | 0.6 | 0.1 | 0.4 | 0.1 |

**Table S2.** d-rGO-B antimicrobial activity at different concentrations in contact with P. aeruginosa and S. epidermidis.

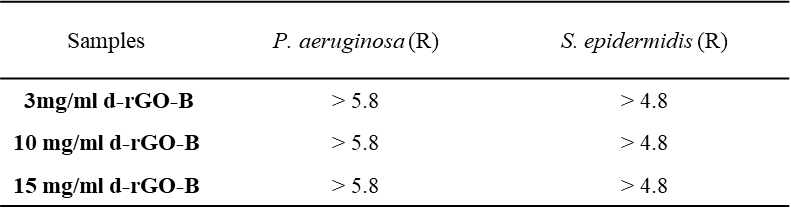

**Table S3*.*** Antimicrobial activity against P. aeruginosa and S. epidermidis of PEOT/PBT- d-rGO-B films (0.1 g/mL) containing different d-rGO concentrations.

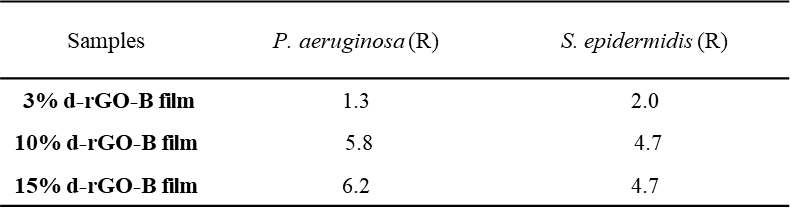
